# Single-cell transcriptomic integrated with machine learning reveals retinal cell-specific biomarkers in diabetic retinopathy

**DOI:** 10.1101/2025.06.03.657753

**Authors:** Sen Lin, Luning Yang, Yiwen Tao, Qi Pan, Tengda Cai, Yunyan Ye, Jianhui Liu, Yang Zhou, Yongqing Shao, Quanyong Yi, Zen Huat Lu, Lie Chen, Gareth McKay, Richard Rankin, Fan Li, Weihua Meng

**Affiliations:** Nottingham Ningbo China Beacons of Excellence Research and Innovation Institute, University of Nottingham Ningbo China, Ningbo, China, 315100; Department of Ophthalmology, Lihuili Hospital affiliated with Ningbo University, Ningbo, China, 315040; Department of Cardiology, Lihuili Hospital affiliated with Ningbo University, Ningbo, China, 315040; Ningbo Institute of Innovation for Combined Medicine and Engineering, The Affiliated Li Huili Hospital, Ningbo University, Ningbo, China, 315201; Department of Ophthalmology, The Affiliated Ningbo Eye Hospital of Wenzhou Medical University, Ningbo, China, 315040; PAPRSB Institute of Health Sciences, Universiti Brunei Darussalam, Bandar Seri, Begawan, Brunei Darussalam, BE1410; Centre for Public Health, Institute of Clinical Science, Queen’s University Belfast, Block B, Royal Hospital, Grosvenor Road, Belfast, UK, BT12 6BA; Centre for English Language Education, University of Nottingham Ningbo China, Ningbo, China, 31510; Eye Center, Zhongshan City People’s Hospital, Zhongshan, Guangdong, China, 528403; Division of Population Health and Genomics, School of Medicine, University of Dundee, Dundee, UK, DD2 4BF; Center for Public Health, Faculty of Medicine, Health and Life Sciences, School of Medicine, Dentistry and Biomedical Sciences, Queen’s University Belfast, Belfast, UK, BT12 6BA

**Keywords:** Single-cell RNA sequencing, Chinese human retina, Machine learning classification, Cell-type-specific biomarkers, Diabetic retinopathy, Cell atlas

## Abstract

Diabetic retinopathy (DR) remains a principal cause of vision impairment worldwide, involved complex retinal cellular pathophysiology that remains incompletely understood. To elucidate cell-type-specific molecular signatures underlying DR, we generated a high-resolution single-cell transcriptomic atlas of 297,121 retinal cells from 20 Chinese donors, including non-diabetic controls (26.4%), diabetic without retinopathy (23.4%) and DR (50.2%). Following rigorous quality control, batch-effect correction, and clustering and annotation, 10 major retinal cell populations were delineated. Differential expression analyses across disease states within each cell type yielded candidate gene sets, which were further refined via a multi-stage machine-learning pipeline combining L1-regularized logistic regression and recursive feature elimination with cross-validation, alongside bootstrap stability selection. Resulting cell-type-specific classifiers achieved high accuracy (79–95%) and AUCs (0.85–0.99) in distinguishing DR disease states. Enrichment analyses implicated immune activation, oxidative stress, neurodegeneration and synaptic dysfunction pathways across multiple cell types in retina. Integrating 567 unique marker genes from all cell types, a general multilayer perceptron classifier achieved 95.31% overall accuracy on held-out test data, demonstrating the translational potential of these signatures for non-diabetic controls, diabetic without retinopathy and DR classification. This high-resolution atlas and the accompanying analytic framework provide a robust computational framework for biomarker discovery, mechanistic insight and targeted intervention strategies in diabetic retinal diseases.

## Introduction

Diabetic Retinopathy (DR) remains one of the most significant complications of diabetes, characterized by complex pathological changes that include retinal inflammation, vascular leakage, and neurodegeneration, leading to irreversible vision impairment in a substantial proportion of diabetic individuals. As one of the foremost causes of preventable blindness worldwide, its progression intricately linked to chronic hyperglycemia in individuals with diabetes [1]. The pathology of DR is characterized by a range of cellular dysfunctions, including retinal inflammation, oxidative stress, and vascular abnormalities, which are not fully understood at the molecular level [2–3]. Early and accurate diagnosis of DR, coupled with a deeper understanding of its underlying molecular mechanisms, is crucial for the development of targeted therapeutic interventions [4]. The complexities of these pathological processes are compounded by the extensive cellular heterogeneity inherent in the retina with its diverse cellular populations and intricate tissue architecture [5]. This complexity necessitates a high-resolution approach capable of distinguishing subtle differences in gene expression within specific retinal cell populations, to unravel the cellular and molecular signatures that underlie diabetic retinal diseases.

Recent advances in single-cell RNA sequencing (scRNA-seq) have revolutionized our ability to study gene expression at the resolution of individual cells, providing a powerful tool to unravel the complexity of retinal tissues [6]. By enabling the profiling of thousands of genes across distinct cellular subpopulations, scRNA-seq offers a deeper insight into the transcriptional landscape of the retina, facilitating the identification of both well-characterized and previously unrecognized cellular signatures in health and disease. When applied to DR, scRNA-seq allows for a more granular understanding of how retinal cells such as retinal ganglion cells, photoreceptors, and glial cells in the disease state, for instance, respond to the metabolic stress. Previous research presented that single-cell analysis identified microglia as a cell type closely associated with lactate metabolism, deepening our understanding of the mechanisms of proliferative diabetic retinopathy [7]. Another study performed scRNA-seq analysis on retinal samples from patients with proliferative DR, identified three key metaprograms, and revealed the role of microglia in the pathogenesis of proliferative diabetic retinopathy [8]. However, the immense volume and complexity of the resulting data present significant challenges for analysis and interpretation, particularly when attempting to integrate information across multiple disease states and cell types.

In this study, we firstly constructed a single-cell transcriptomic atlas of the Chinese human retina, utilizing scRNA-seq data derived from a cohort of both living donors and postmortem specimens. The dataset encompassed a wide array of retinal cell types, including neurons, glial cells, and photoreceptors, providing a holistic view of retinal cellular heterogeneity. By integrating high-quality data from these samples, we systematically characterized the gene expression profiles of distinct retinal cell populations, with a particular focus on identifying alterations associated with diabetic and diabetic retinopathy conditions. To identify the most relevant gene markers for DR, we employed a robust computational framework that combines machine learning techniques, including L1-regularized logistic regression and recursive feature elimination with cross-validation (RFECV), to systematically select genes that can distinguish between non-diabetic and diabetic retinal states. Furthermore, we developed neural network-based models to classify disease states based on these gene signatures, achieving high accuracy in distinguishing diabetic and non-diabetic retinal samples. The integration of these advanced methodologies facilitates a deeper understanding of the retinal cellular architecture in DR, uncovering the molecular mechanisms that contribute to retinal change in diabetic patients. Ultimately, this study enabled the detection of subtle transcriptional changes that may not be apparent when analyzing bulk tissue, thus revealing the intricate molecular mechanisms that differentiate retinal cells in different disease states.

## Materials and Methods

### Human retinal tissue collection and processing

Ethical approval for the collection of human retinal tissue was formally granted by the ethics committees of the University of Nottingham Ningbo China, the Ningbo Eye Hospital, and the Ningbo Medical Centre Lihuili Hospital. 20 human retinal tissue samples were collected June 2023 and March 2025, with 18 sourced from the Ningbo Eye Hospital and 2 from the Lihuili Hospital. Written informed consent was obtained from living donors prior to tissue acquisition. Retinal tissues included 7 bilateral pairs and 1 unilateral specimen from deceased individuals, alongside 5 unilateral surgical enucleation-derived samples (detailed in Table 1). In this study, fresh retinal tissues were defined as either specimens harvested from living donors within 10 minutes after surgical enucleation completion or postmortem specimens retrieved within 6 hours postmortem under strictly monitored temperature conditions.

**Table 1:** Donor sample information.

| Sample | Eye | Sex | Age | Diabetic Status | Eye Removal/Death Reason |
| --- | --- | --- | --- | --- | --- |
| 01 | Left | M | 55 | Non | Eyeball atrophy |
| 02 | Right | M | 71 | Non | Corneal ulcer |
| 03 | Left | F | 50 | Non | Eyeball atrophy |
| 04 | Right | F | 68 | Non | Corneal perforation |
| 05 | Left | M | 34 | Non | Cerebral hemorrhage |
| 06 | Left | F | 92 | Non | Natural death |
| 07 | Right | F | 92 | Non | Natural death |
| 08 | Left | M | 76 | DM | Corneal staphyloma |
| 09 | Left | F | 55 | DM | SCT-H-PON |
| 10 | Right | F | 55 | DM | SCT-H-PON |
| 11 | Left | M | 87 | DM | Liver cancer |
| 12 | Right | M | 87 | DM | Liver cancer |
| 13 | Left | M | 55 | DR | Brain stem hemorrhage |
| 14 | Right | M | 55 | DR | Brain stem hemorrhage |
| 15 | Left | M | 35 | DR | Traumatic shock |
| 16 | Right | M | 35 | DR | Traumatic shock |
| 17 | Left | M | 84 | DR | Prostate malignancy |
| 18 | Right | M | 84 | DR | Prostate malignancy |
| 19 | Left | M | 69 | DR | Lung cancer |
| 20 | Right | M | 69 | DR | Lung cancer |

DM: Samples with diabetes. DR: Samples with diabetic retinopathy. non: Samples without diabetes. SCT-H-PON: Severe craniocerebral trauma with hemorrhage in the periphery of the optic nerve. As retinal tissues were acquired during surgical procedures or post-mortem and detailed fundus imaging was not available, the specific duration of diabetic retinopathy could not be determined for these DR cases. 20 samples were from 13 donors, with sample 06 and 07, sample 09 and 10, sample 11 and 12, sample 13 and 14, sample 15 and 16, sample 17 and 18, sample 19 and 20 from the left and right eyes of the same donor.

### Single-cell RNA sequencing technique

Human retinal tissues were dissociated for scRNA-seq underwent standardized dissociation protocols utilizing the Worthington Papain Dissociation (catalog no. LK003150-1bx, Worthington Biochemical Corporation). Fresh retinal tissues were chilled in ice-cooled Petri dishes and mechanically sectioned into 2-4 mm fragments. Enzymatic digestion commenced with immersion in RPMI-1640 medium supplemented with Miltenyi Biotec enzymatic mixture (37°C, 15 min), followed by mechanical processing in gentleMACS C Tubes with supplemented additional RPMI/enzyme mix. Post-digestion suspensions underwent sequential filtration through 70 µm cell strainers and centrifugation (300×g, 4°C, 10 min). Erythrocyte depletion was achieved through 10 times volumes of 1× volume Red Blood Cell Lysis Solution (Miltenyi Biotec, Cat#130-094-183; 4°C, 10 min), followed by 1x PBS/0.04% BSA resuspension. Cell viability (>85%) and concentration (700-1200 cells/μL) were quantified via CountStar prior to library construction. Chromium Single Cell 3’ Library & Gel Bead Kit (10X Genomics, PN1000268) was employed for 10,000 viable cells per sample, with sequenced on the Illumina platform after stringent quality control: exclusion of reads containing >3 ambiguous bases, >20% low-quality bases (quality rate <5), or adapter contaminants.

### Generation of single-cell RNA sequencing data

Raw sequencing reads underwent demultiplexing and reference alignment (GRCh38-2020-A assembly) through the 10X Genomics Cell Ranger pipeline (v9.0.0) employing standard pipeline configurations (https://www.10xgenomics.com/support/software/cell-ranger). This systematic processing pipeline executed four core functions: sequence alignment, quality filtering, cellular barcode enumeration, and unique molecular identifier (UMI) quantification. The algorithm leveraged Chromium-specific cellular barcodes to construct feature-barcode matrices that powered downstream computational workflows encompassing dimensionality reduction, cluster identification, and transcriptome profiling through analytical frameworks.

### Quality control and doublets removal

Processed sequencing outputs from the 10X Genomics Cell Ranger pipeline (cellular barcodes, gene feature identifiers, and count matrices) underwent rigorous quality control through some filtering methods. High-quality cell data was operationally defined by three exclusion criteria: minimum 100 expressed genes per cell, greater than 400 total UMI counts per cell, and less than 10% mitochondrial gene content. Substandard cell data were systematically excluded. Subsequent data refinement incorporated computational doublet elimination via Scrublet (v0.2.3) [9], implementing a Python-based k-nearest neighbor algorithm that discriminates simulated artificial doublets from empirical transcriptomic profiles through comparative manifold learning.

### Batch effect and data integration

Post-quality control process yielded 20 high-fidelity datasets for downstream analysis. To address batch effects, a critical confounding factor in multi-sample integration that can precipitate misclassification or distort results in subsequent analytical workflows, we implemented Batch Balanced K Nearest Neighbours (BBKNN) (v1.6.0) [10]. For each cell, BBKNN identifies the k nearest neighbors within each individual batch and then aggregates these batch-specific neighbor lists into a unified neighbor graph. By constructing balanced cellular neighborhoods across batches while preserving intrinsic biological variation, BBKNN aligns phenotypically similar cells across disparate samples, thereby effectively mitigates technical biases arising from samples.

### Data normalization, transformation and scaling

ScRNA-seq data preprocessing was conducted via Scanpy (v1.11.1) [11] to address technical confounders including sequencing depth variance, capture efficiency and library preparation heterogeneity. Raw scRNA-seq data often displayed pronounced skewness, which can impair downstream steps including dimensionality reduction, clustering, and differential expression testing. Raw counts were first normalized and then log-transformed (log1p) to stabilize variance and promote comparability of gene expression levels across cells. Following normalization, the log1p transformation and scaling, all cells were distributed on a common numerical scale, thereby minimizing technical biases, achieving a more symmetric and even data distribution, and ensuring that subsequent analyses more faithfully represent intrinsic biological variation.

### Data dimensionality reduction and cell clustering

Single-cell transcriptomic datasets typically measure thousands of genes, many of which show minimal variation and do not contribute to distinguishing cell types but increase dimensionality while introducing stochastic noise. To optimize feature selection, we implemented a variance stabilization protocol retaining the top 5,000 highly variable genes. These feature selection reduced dimensionality while preserving key information of cell state and identity that exhibited cell type-discriminatory potential. Principal Component Analysis (PCA) was subsequently employed for dimensional reduction to the filtered variable gene expression matrix to project high-dimensional data into a lower-dimensional space. This linear transformation effectively maximized variance capture and segregated biologically informative signals and meaningful differences from technical artifacts and background noise.

To partition cells into distinct groups, we used the Leiden graph-based clustering algorithm to delineate transcriptionally distinct cell subpopulations [12]. A k-nearest neighbors (KNN) graph for each candidate resolution value between 0.02 and 2.00 was constructed, and the algorithm iteratively identified clustering that reflect biologically coherent cell populations. For each resolution setting, we generated uniform manifold approximation and projection (UMAP) to project the clustered cells into two-dimensional space, enabling intuitive visualization and interpretation of cluster-specific expression patterns.

### Cell type annotation

Cell type annotation was conducted manually to delineate discrete subpopulations through integrative analysis of dimensionality-reduced embeddings and established marker genes. UMAP afforded a robust two-dimensional projection of individual cells, facilitating the visual discrimination of clusters corresponding to distinct cell types or states. By integrating canonical marker gene expression profiles with existing biological knowledge, each cluster was assigned to a specific cell type, ensuring accurate interpretation of cellular identities.

### Differential expression analysis

Differential expression analysis was systematically performed to identify genes whose expression levels differ significantly between samples with diabetes (DM), diabetic retinopathy (DR), and non-diabetic controls (NON). Three comparative cohorts were interrogated: DM versus NON, DR versus NON, and DR versus DM. It applied a nonparametric Wilcoxon rank-sum test to each gene, adjusted p-values via the Benjamini–Hochberg procedure to control the false discovery rate (FDR), filtered for robust effect sizes (logL fold-change ≥ 1) and significance (adjusted p < 0.05), prioritized genes exhibiting both statistical robustness and biological relevance, and exported the resulting differential expression gene lists for downstream interpretation.

### Important genes identification

After cell type annotation, cells were categorized into distinct types based on established retinal cellular atlas. Subsequently, differential expression analyses were conducted within each cell type individually, thereby isolating the gene expression variations related to each disease state. For each cell type in the retina, we designed a rigorous, multiLstage machineLlearning framework to identify important genes for distinguishing disease states. The top 50 genes with the highest absolute log2-fold changes from each comparison resulting differential expression gene list were selected and merged into a unique gene feature set after deduplication. The gene expression matrix of unique gene feature set the were extracted from the whole gene expression matrix and normalized using z-score transformation. The resulting gene expression matrix was then split into 70 % training and 30 % testing cohorts under stratified sampling to preserve disease-state class proportions, and all features were centered and scaled to unit variance to satisfy the assumptions of linear models.

Feature selection employed a two-stage approach: first, L1-regularized logistic regression with 5-fold cross-validation grid search (GridSearchCV) optimized the regularization parameter C (0.01-10 range) to induce sparsity and identify the most discriminative genes within a oneLversusLrest classification framework [13]. Subsequently, to further refine the feature space, recursive feature elimination with internal crossLvalidation (RFECV) was applied around the tuned logistic model, iteratively removing the least informative genes until peak predictive performance was reached [14]. Recognizing selection variability, we then conducted a stability analysis by repeating recursive feature elimination (RFE) across 20 bootstrap replicates; genes selected in more than half of these runs were designated as robust markers. This multiLstage machineLlearning framework not only ensures that the selected gene features can fully express the differences between disease states while avoiding overfitting, but also provide more stable and interpretable resulting gene features for subsequent analysis.

Finally, we assessed model performance on the heldLout test set through precision, recall, and F1Lscore metrics, complemented by oneLversusLrest ROC curves and corresponding AUC values for each disease-state class. This comprehensive pipeline from differential expression filtering through sparse modeling, recursive selection, and stability validation ensured both parsimony and reproducibility in the identification of disease states–specific important genes for each cell type. This dual regularization strategy (L1 penalty + RFECV) balanced model sparsity with discriminative power while stability selection enhances biological interpretability of marker genes. Meanwhile, this methodology ensures that observed differential gene expression reflects true biological differences rather than artifacts introduced by cell type composition disparities. Employing this cell type-specific analysis framework enhances the reliability of identifying disease-associated gene and facilitates a more accurate understanding of the molecular underpinnings of retinal pathophysiology.

### Neural Network-Based Classification

To further verify that important genes can be used to identify retinal cells in disease states, we merged the important genes of each retinal cell type to a gene list. The gene expression matrix of this gene list was extracted and partitioned using stratified sampling (70% training, 30% testing) to maintain disease-state class distribution parity, followed by z-score normalization to standardize feature scales. We then trained a two–hiddenLlayer multilayer perceptron (MLP) classifier using scikit-learn (v1.5.1), with hidden layer sizes of 100 neurons each, and rectified linear unit (ReLU) activation functions. The model employed adaptive learning rate optimization via the Adam solver, L2 regularization, and early stopping based on convergence thresholds (training proceeded for up to 200 epochs with verbose logging to monitor loss reduction). Training stability was ensured through batch-wise gradient updates and fixed randomization. Model performance was quantified using multiclass accuracy, precision-recall metrics, and a confusion matrix. Visual validation included a heatmap representation of class prediction distributions, annotated with absolute counts. All evaluation metrics (classification report, accuracy score, and confusion matrix) were systematically archived in standardized output files to ensure reproducibility. This architecture leverages non-linear decision boundaries to capture complex gene expression patterns associated with diabetic disease progression [15].

### Enrichment analysis

Gene Ontology (GO) and Kyoto Encyclopedia of Genes and Genomes (KEGG) pathway enrichment analyses were performed to elucidate the biological functions and pathways associated with the important gene lists, leveraging the clusterProfiler R package (v4.12.6) [16–17]. GO and KEGG enrichment analysis facilitates the interpretation of gene clusters by associating them with predefined biological terms. GO enrichment analysis was performed on the Biological Process (BP) ontology to identify over-represented GO terms among the important gene lists. Statistical significance was determined using the Benjamini-Hochberg method to adjust p-values, with a threshold of 0.05. KEGG enrichment analysis maps genes to known pathways to identify biological processes that are significantly associated with the important gene lists. Biological pathways were considered significantly enriched if they had an adjusted p-value less than 0.05. These enrichment analyses provided insights into the biological mechanisms underlying the observed gene expression changes, facilitating a deeper understanding of the functional implications of the important gene lists in the context of cellular biological processes and disease-relevant metabolic pathways.

## Results

### Construction of human retinal scRNA-Seq cell atlas

We established a high-resolution retinal cellular atlas through systematic integration of 20 retinal samples obtained from Chinese living donors and postmortem specimens. Following stringent quality-controlled preprocessing, we employed UMAP to visualize cellular heterogeneity across all samples (Figure 1A and 1B). This analysis revealed 10 distinct retinal cell populations, each characterized by specific marker genes: amacrine cells (ACs): expressing *TFAP2B*, *GAD1*, and *SLC32A1*; astrocytes: identified by *GFAP*; bipolar cells (BCs): marked by *GRM6* and *GRIK1*; cone photoreceptors: identified by *ARR3*, *GNAT2*, and *GNGT2*; horizontal cells (HCs): expressing *LHX1* and *TNR*; microglia: identified by *CD74* and *TYROBP*; müller glial cells (MGCs): characterized by *RLBP1*, *RGR*, and *DKK3*; retinal ganglion cells (RGCs): marked by *RBPMS* and *SLC17A6*; rod photoreceptors: characterized by *RHO*, *PDE6G*, *GNGT1*, and *NRL*; T cells: characterized by *CD69*, *CD52*, and *CD3D* (Figure 1C and 1D) [18].

**Figure 1:**
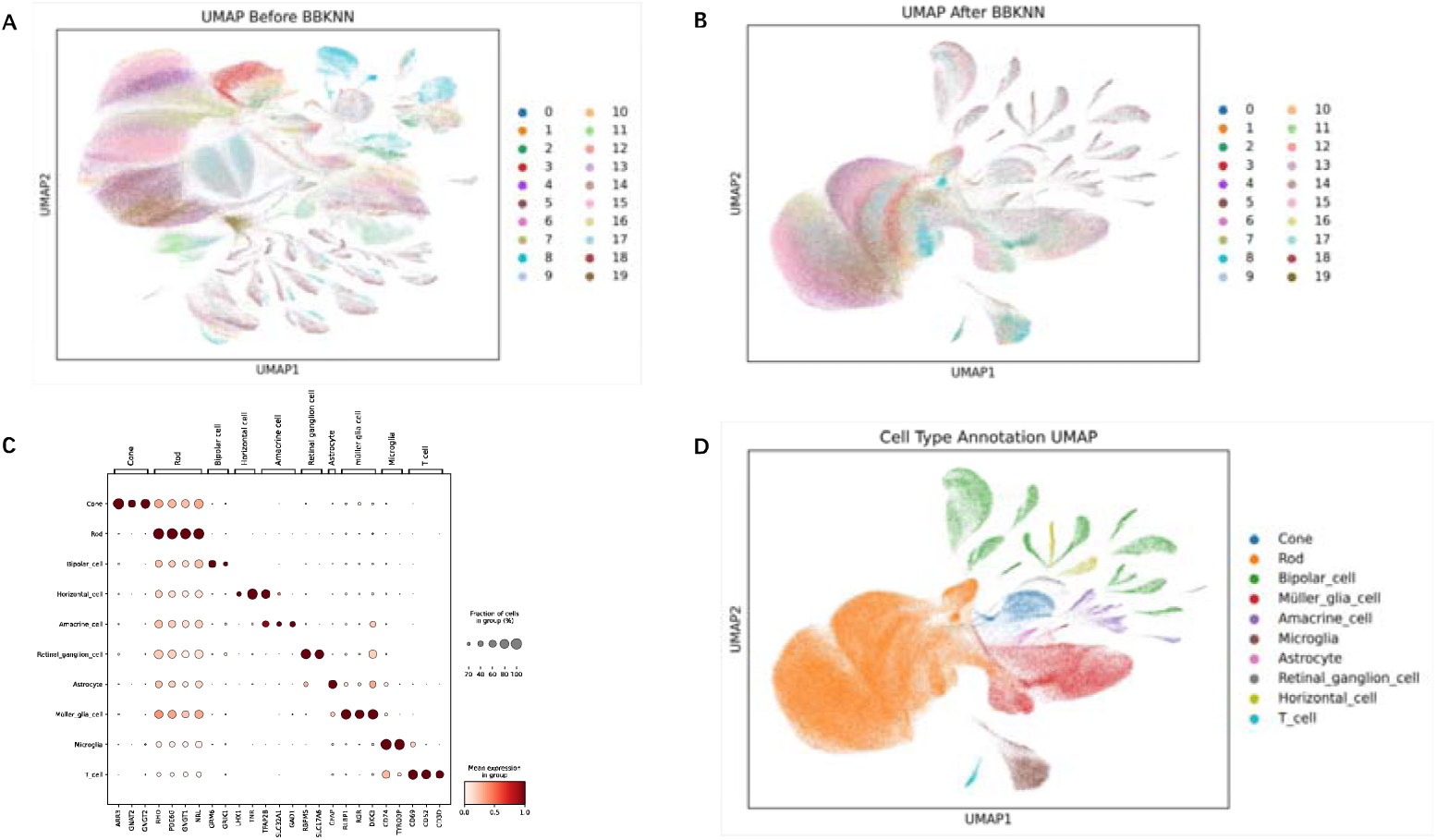
batch effect removal and cell clustering and annotation (A) UMAP projection of integrated scRNA-seq data from 20 Chinese human retinal samples, with each point representing a single cell. The observed uneven cell distribution reflects batch effects across the different samples. (B) UMAP projection of integrated scRNA-seq data from 20 Chinese human retinal samples following batchLeffect correction via Batch Balanced K Nearest Neighbors (BBKNN). PostLBBKNN, cell distribution is independent of sample origin. (C) Dot plot illustrated the annotation of each cluster in retinal samples based on established markers for major cell types, alongside the selection of genes that distinguish individual clusters. Each row represents a specific major cell type, while each column denotes the expression level of a key marker gene. The top annotation highlights the principal marker genes employed for cellLtype identification. (D) UMAP projection of cell clusters annotated according to the major human retinal cell types.

This atlas provided an in-depth characterization of the cellular diversity within the Chinese human retina, offering valuable insights into its complex cellular architecture. The total number of preprocessed cells for downstream analysis was 297,121 from 20 Chinese donor samples (Figure 2B). The cells in DR condition constituted the majority subset (the number is 149,116; accounting for 50.2%), followed by the cells in DM condition (the number is 69,524; accounting for 23.4%) and non-diabetic controls (the number is 78,481; accounting for 26.4%).

**Figure 2:**
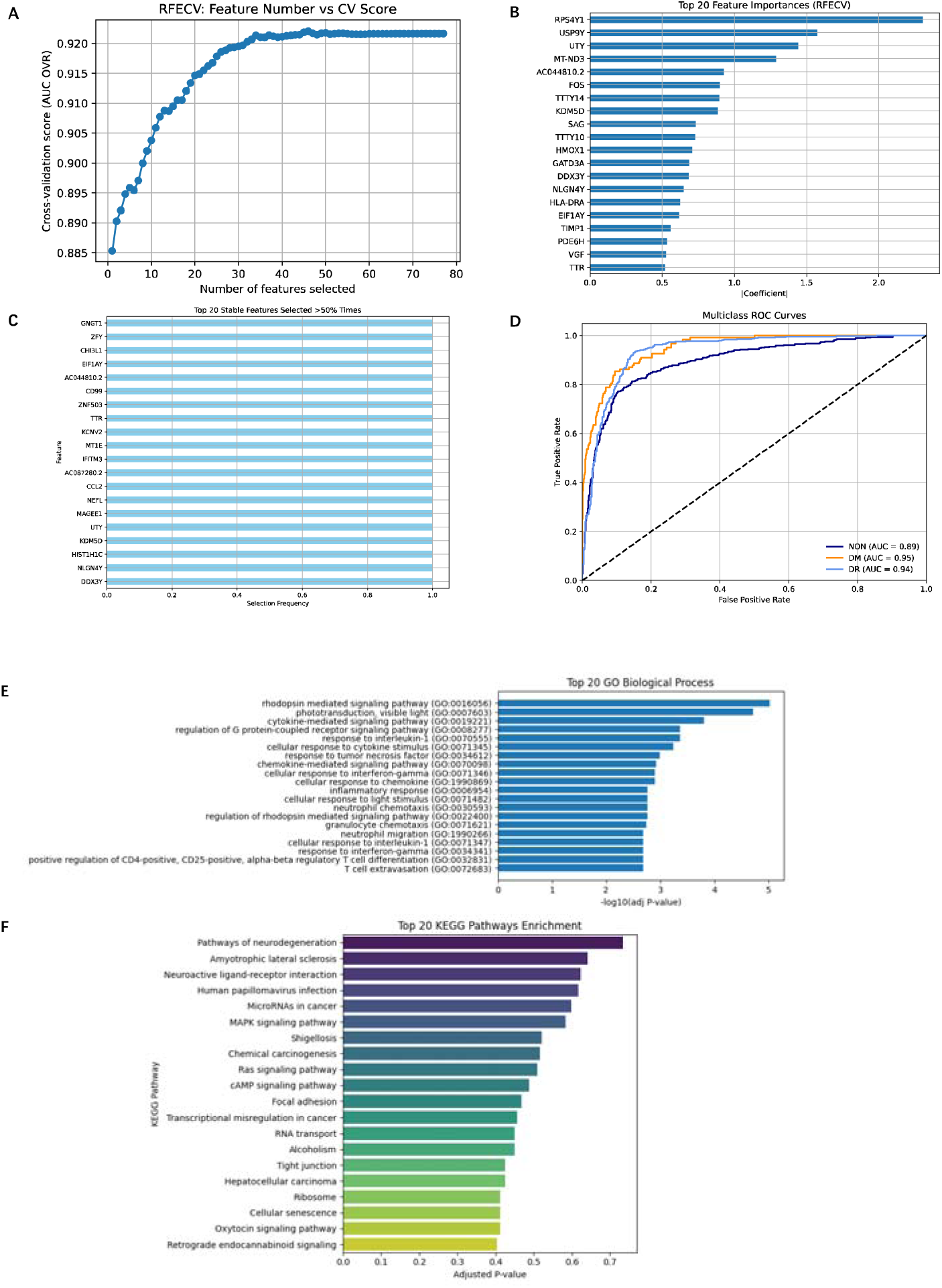
Comprehensive analysis of Chinese human retinal ACs type (A) RFECVLderived feature selection curve: RFECV was performed using an L1-penalized logistic regression classifier under a one-versus-rest scheme. The plot illustrated the relationship between the number of selected features and the corresponding mean cross-validated performance metric (AUC One-vs-Rest). The observed unimodal trend indicated that model performance is maximized with a relatively small feature subset. The optimal feature subset size was therefore chosen for downstream modeling. (B) To assess the relative importance of the features selected by RFECV, we plotted the top 20 features based on the sum of the absolute values of the coefficients across all classes in the logistic regression model. The plot showed the features arranged in descending order of importance. The horizontal bars represent the magnitude of the coefficients, with longer bars indicating greater contribution to the model’s classification performance. Notably, genes such as *RPS4Y1*, *USP9Y*, and *UTY* exhibit the highest feature importances, suggesting their critical roles in distinguishing between the different classes (healthy, diabetes, diabetic retinopathy) in this study. (C) To evaluate the robustness of the feature selection process, we performed stability selection by repeating RFE 20 times with different random seeds. Each run selected a fixed number of features, as determined previously by RFECV. For each gene, we calculated the proportion of times it was selected across the 20 repetitions. The plot displayed the top 20 most frequently selected features among those chosen in over 50% of the runs. Higher selection frequency indicates greater stability and reliability of the feature in contributing to classification performance. Genes such as *GNGT1*, *ZFY*, and *CHI3L1* were consistently retained, suggesting their strong and reproducible discriminative power across subsamples. (D) To evaluate the classification performance of the final model, ROC curves were constructed for each class (NON, DM, DR) using a one-versus-rest approach. The plot displayed the true positive rate plotted against the false positive rate for each label. The colored curves represent model performance for each class, with the diagonal dashed line indicating the performance of a random classifier (AUC = 0.5). The area under the curve (AUC) values, reported in the legend, quantify the model’s ability to discriminate each class. All three classes exhibit AUCs substantially greater than 0.5, indicating strong predictive power and reliable differentiation between healthy, diabetic, and retinopathic states. (E) Bar plot presented the top 20 enriched GO terms for the important and stable gene list. Rows represented each GO term that these genes were involved in. The horizontal bars represent the p-value, the longer bars indicating the stronger the correlation. (F) Bar plot presented the top 20 enriched KEGG terms for the important and stable gene list. Rows represented each KEGG term that these genes were involved in. The horizontal bars represent the adjusted p-value, the longer bars indicating the stronger the correlation.

### Machine□learning Based Analysis of Human Retinal Cell-Type Specific Disease-associated Genes

To enhance the precision of differential gene expression analyses across disease states DR, DM, and non-diabetic controls (NON) retinal samples, and to mitigate confounding effects arising from cellular type heterogeneity, we implemented a cell type-specific stratification approach. For each human retinal cell type, the multiLstage machineLlearning frameworks (L1-regularized logistic regression and RFECV) were sequentially performed to dynamically and systematically identify, evaluate and select the most discriminative gene set for disease states. Then we conducted RFE stability selection by repeating RFE across 20 bootstrap replicates to ensure the stability and robustness of the selected gene sets. GO and KEGG enrichment analysis as pivotal tools in bioinformatics was employed, aiding in elucidating the functional roles of genes within complex biological systems and understanding their involvement in various physiological and pathological processes.

### Amacrine Cells

Aimed to determine the optimal number of selected genes, we applied RFECV using an L1-regularized logistic regression estimator under a one-versus-rest scheme (Figure 2A). The number of selected important genes to distinguish between DR, DM and non-diabetic controls ACs were 55 through RFECV model (Supplementary Table 1). Figure 2B illustrates the top 20 genes with the highest feature weights in the model. Subsequent RFE stability selection results remained the same for these 55 genes. Figure 2C demonstrates that the top 20 features are the most reliable across many slightly different fits. To further validate these selected important genes, we trained a one-versus-rest logistic regression classifier only using these selected important genes as features. The classifier achieved a high degree of accuracy (85%), and the multiclass ROC curves yielded AUCs of AUC_NON (0.89), AUC_DM (0.95), and AUC_DR (0.94), demonstrating that the selected genes discriminated the disease sates of samples with high accuracy (Figure 2D). The selected genes identified in ACs are implicated in various biological processes that contribute to the disease’s pathogenesis. Genes such as *CHI3L1* and *CCL2* are involved in inflammatory pathways, linked to neuroinflammatory conditions and contributed to leukocyte infiltration and inflammation [19–20]. Genes like *GNGT1*, *KCNV2*, and *NEFL* are crucial for neuronal signaling [21]. Gene *MT1E* is part of the metallothionein family, which plays a role in metal ion homeostasis and protection against oxidative stress that is a significant factor in DR progression [22]. Genes such as *KDM5D*, *UTY*, and *DDX3Y* are involved in chromatin remodeling and transcriptional regulation, which may influence gene expression patterns in response to hyperglycemic conditions, affecting cell survival and function [23–25].

We performed GO enrichment analysis on these selected genes to list the biological process items involved such as “rhodopsin mediated signaling pathway”, “phototransduction, visible light”, and “cytokine-mediated signaling pathway”. (Figure 2E). KEGG enrichment analysis elucidated the functional roles of these genes within complex biological systems, physiological and pathological processes, for instance “pathways of neurodegeneration”, “amyotrophic lateral sclerosis”, and “neuroactive ligand-receptor interaction”. (Figure 2F).

### Bipolar Cells

Using the same multi-stage machine learning framework (L1-regularized logistic regression + RFECV with one-versus-rest) and RFE stability selection over 20 bootstrap replicates (Figure 3A), RFECV identified 96 key genes for discriminating DR, DM, and non-diabetic controls in BCs (Supplementary Table 1). The top 20 features by absolute weight are shown in Figure 3B, and stability selection confirmed their consistent selection across replicates. RFE stability plots (Figure 3C) further underscore the robustness of this gene set. A one-versus-rest logistic regression classifier trained solely on these 96 genes achieved 83% accuracy, with multiclass ROC AUCs of 0.89 (NON), 0.93 (DM), and 0.97 (DR) (Figure 3D). The genes identified in BCs were associated with several biological processes, including immune response, inflammation, oxidative stress, and phototransduction. Genes such as *C1QB*, *CD163*, *CCL2*, *CCL3L1*, *CXCL8* are involved in immune and inflammatory pathways [26]. Gene *FCER1G* is associated with immune cell activation and has been linked to macrophage infiltration in various diseases [27]. Gene *S100A8* is a calcium-binding protein that participates in inflammatory processes and has been implicated in diabetic complications [28]. Gene *CYBA* encodes oxidase complex, leads to oxidative stress, a key factor in DR pathogenesis [29]. Genes *PDE6A* and *PDE6G* are crucial in the phototransduction cascade and dysfunction in them may affect visual signal transduction [30]. The expression of gene *HIST1H4C* are related to hyperglycemia [31].

**Figure 3:**
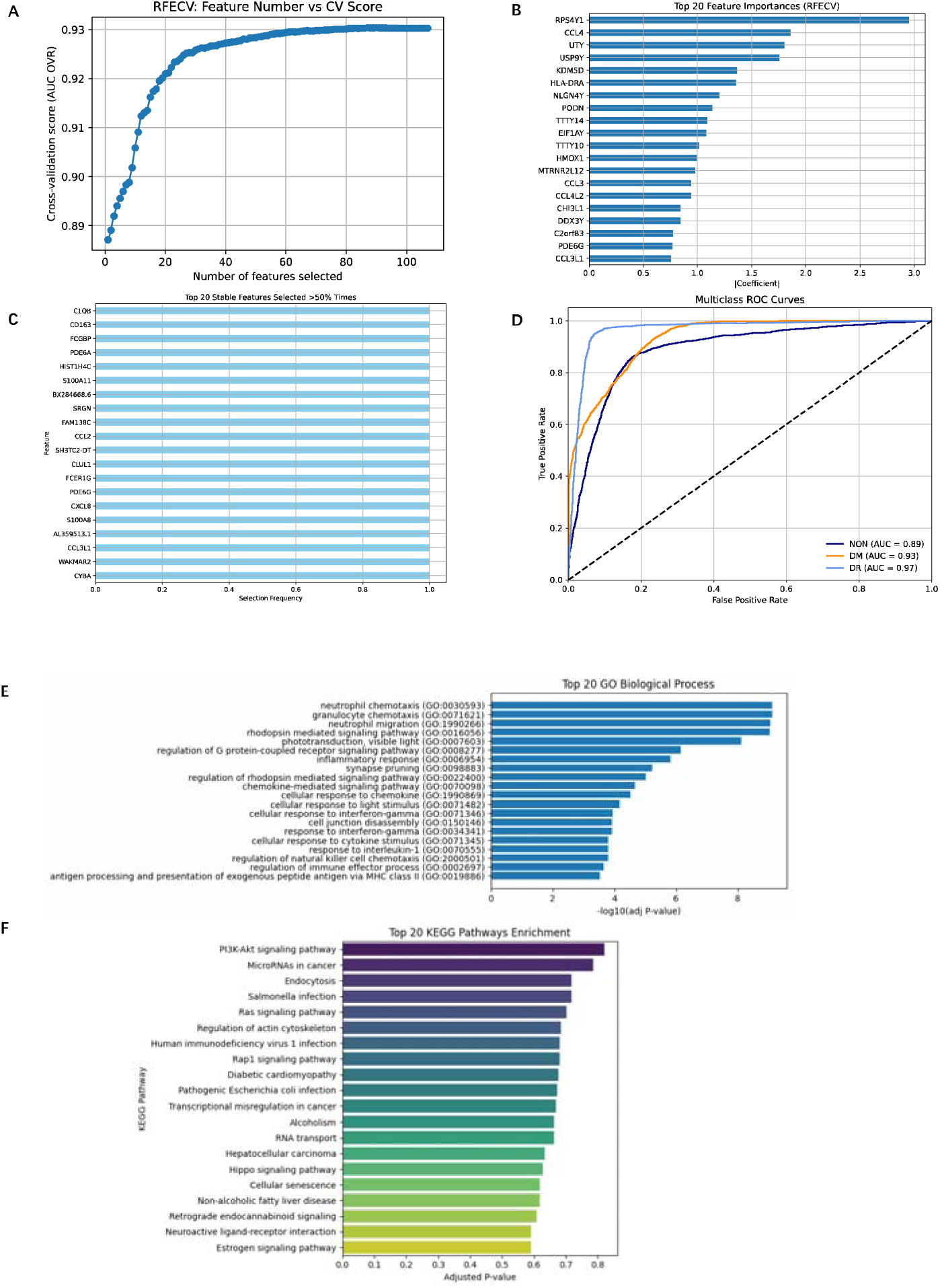
Comprehensive analysis of Chinese human retinal BCs type (A) RFECVLderived feature selection curve: RFECV was performed using an L1-penalized logistic regression classifier under a one-versus-rest scheme. The plot illustrated the relationship between the number of selected features and the corresponding mean cross-validated performance metric (AUC One-vs-Rest). The observed unimodal trend indicated that model performance is maximized with a relatively small feature subset. The optimal feature subset size was therefore chosen for downstream modeling. (B) To assess the relative importance of the features selected by RFECV, we plotted the top 20 features based on the sum of the absolute values of the coefficients across all classes in the logistic regression model. The plot showed the features arranged in descending order of importance. The horizontal bars represent the magnitude of the coefficients, with longer bars indicating greater contribution to the model’s classification performance. Notably, genes such as *RPS4Y1*, *CCL4*, and *UTY* exhibit the highest feature importances, suggesting their critical roles in distinguishing between the different classes (healthy, diabetes, diabetic retinopathy) in this study. (C) To evaluate the robustness of the feature selection process, we performed stability selection by repeating RFE 20 times with different random seeds. Each run selected a fixed number of features, as determined previously by RFECV. For each gene, we calculated the proportion of times it was selected across the 20 repetitions. The plot displayed the top 20 most frequently selected features among those chosen in over 50% of the runs. Higher selection frequency indicates greater stability and reliability of the feature in contributing to classification performance. Genes such as *C1QB*, *CD163*, and *FCGBP* were consistently retained, suggesting their strong and reproducible discriminative power across subsamples. (D) To evaluate the classification performance of the final model, ROC curves were constructed for each class (NON, DM, DR) using a one-versus-rest approach. The plot displayed the true positive rate plotted against the false positive rate for each label. The colored curves represent model performance for each class, with the diagonal dashed line indicating the performance of a random classifier (AUC = 0.5). The area under the curve (AUC) values, reported in the legend, quantify the model’s ability to discriminate each class. All three classes exhibit AUCs substantially greater than 0.5, indicating strong predictive power and reliable differentiation between healthy, diabetic, and retinopathic states. (E) Bar plot presented the top 20 enriched GO terms for the important and stable gene list. Rows represented each GO term that these genes were involved in. The horizontal bars represent the p-value, the longer bars indicating the stronger the correlation. (F) Bar plot presented the top 20 enriched KEGG terms for the important and stable gene list. Rows represented each KEGG term that these genes were involved in. The horizontal bars represent the adjusted p-value, the longer bars indicating the stronger the correlation.

GO enrichment revealed significant overrepresentation of “neutrophil chemotaxis”, “granulocyte chemotaxis”, and “neutrophil migration” (Figure 3E). KEGG analysis highlighted “PI3K-Akt signaling pathway”, “microRNAs in cancer”, and “endocytosis” (Figure 3F), suggesting that the related biological process items altered in diabetic conditions.

### Retinal Ganglion Cells

Applying the same framework, RFECV selected 26 discriminative genes in RGCs (Figure 4A and Supplementary Table 1). The foremost 20 genes by coefficient magnitude are displayed in Figure 4B, with RFE stability selection also confirming the stable genes reproducibility across bootstraps (Figure 4C). A classifier based on these 26 RFE stability selected genes yielded 93% accuracy and ROC AUCs of 0.97 (NON), 0.99 (DM), and 0.97 (DR) (Figure 4D). The stable gene feature selection reinforces the model’s reliability. Various biological processes contribute to the pathogenesis of DR by affecting RGCs survival and function. Gene *B2M* have been associated with inflammatory responses in diabetic conditions [32]. Gene *GATD3A* is involved in mitochondrial function and may play a role in managing oxidative stress within RGCs, which is a significant factor in DR progression [33]. Genes such as *LINC01115*, *LINC01933*, *LINC01303*, and *UNC5B-AS1* are involved in the regulation of gene expression at the transcriptional and post-transcriptional levels, which may modulate pathways related to inflammation, apoptosis, and oxidative stress in RGCs during DR [34].

**Figure 4:**
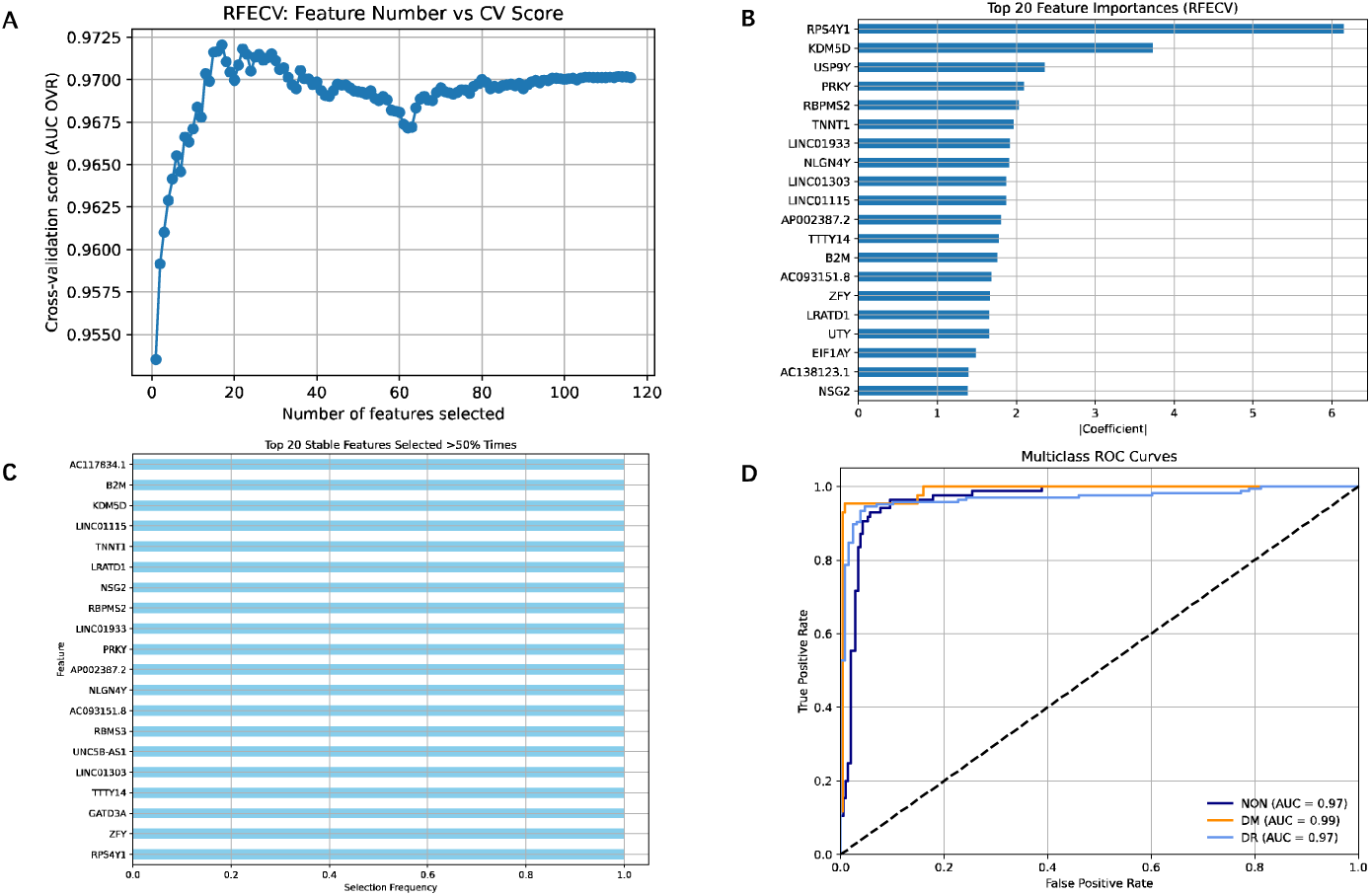
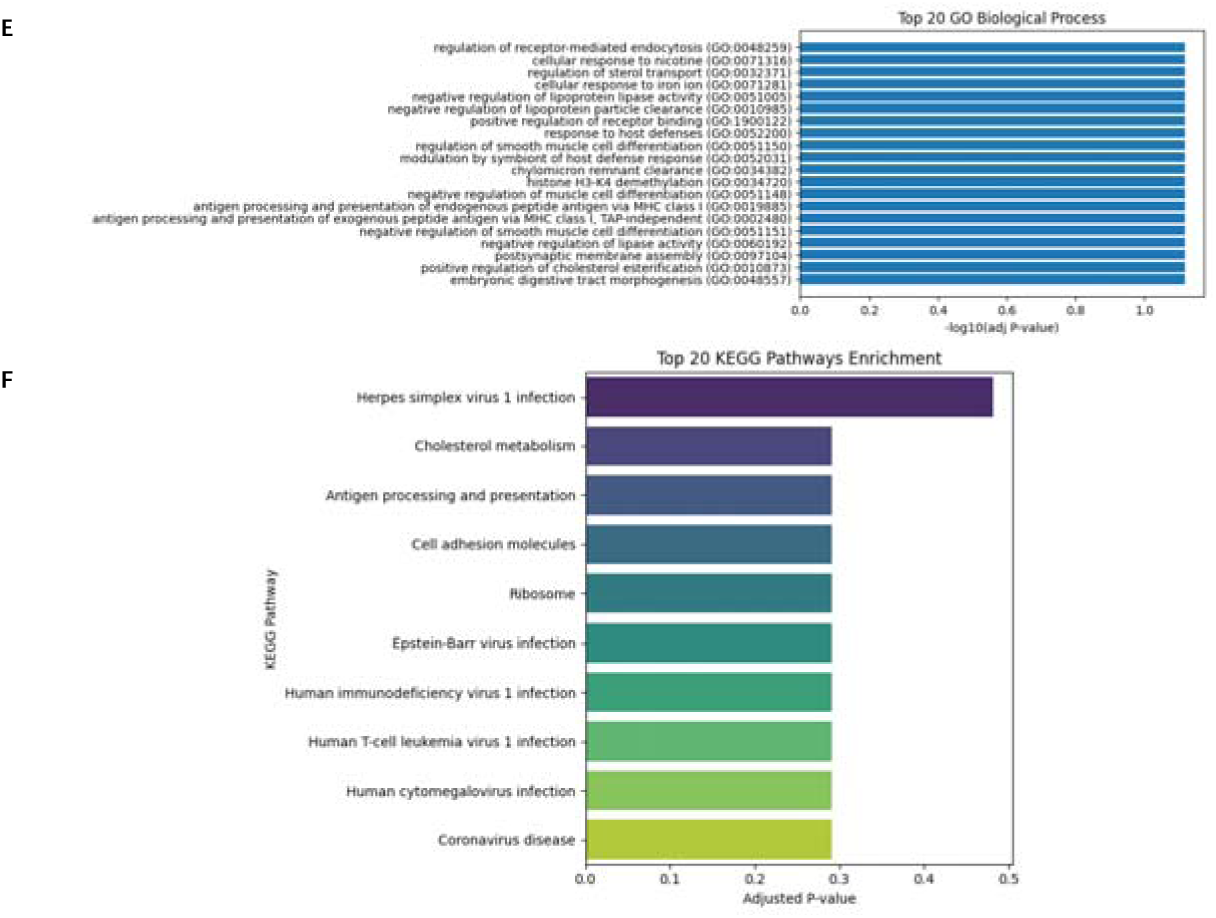
Comprehensive analysis of Chinese human retinal RGCs type (A) RFECVLderived feature selection curve: RFECV was performed using an L1-penalized logistic regression classifier under a one-versus-rest scheme. The plot illustrated the relationship between the number of selected features and the corresponding mean cross-validated performance metric (AUC One-vs-Rest). The observed unimodal trend indicated that model performance is maximized with a relatively small feature subset. The optimal feature subset size was therefore chosen for downstream modeling. (B) To assess the relative importance of the features selected by RFECV, we plotted the top 20 features based on the sum of the absolute values of the coefficients across all classes in the logistic regression model. The plot showed the features arranged in descending order of importance. The horizontal bars represent the magnitude of the coefficients, with longer bars indicating greater contribution to the model’s classification performance. Notably, genes such as *RPS4Y1*, *KDM5D*, and *USP9Y* exhibit the highest feature importances, suggesting their critical roles in distinguishing between the different classes (healthy, diabetes, diabetic retinopathy) in this study. (C) To evaluate the robustness of the feature selection process, we performed stability selection by repeating RFE 20 times with different random seeds. Each run selected a fixed number of features, as determined previously by RFECV. For each gene, we calculated the proportion of times it was selected across the 20 repetitions. The plot displayed the top 20 most frequently selected features among those chosen in over 50% of the runs. Higher selection frequency indicates greater stability and reliability of the feature in contributing to classification performance. Genes such as *AC117834.1*, *B2M*, and *KDM5D* were consistently retained, suggesting their strong and reproducible discriminative power across subsamples. (D) To evaluate the classification performance of the final model, ROC curves were constructed for each class (NON, DM, DR) using a one-versus-rest approach. The plot displayed the true positive rate plotted against the false positive rate for each label. The colored curves represent model performance for each class, with the diagonal dashed line indicating the performance of a random classifier (AUC = 0.5). The area under the curve (AUC) values, reported in the legend, quantify the model’s ability to discriminate each class. All three classes exhibit AUCs substantially greater than 0.5, indicating strong predictive power and reliable differentiation between healthy, diabetic, and retinopathic states. (E) Bar plot presented the top 20 enriched GO terms for the important and stable gene list. Rows represented each GO term that these genes were involved in. The horizontal bars represent the p-value, the longer bars indicating the stronger the correlation. (F) Bar plot presented the top 20 enriched KEGG terms for the important and stable gene list. Rows represented each KEGG term that these genes were involved in. The horizontal bars represent the adjusted p-value, the longer bars indicating the stronger the correlation.

Enriched GO terms included “regulation of receptor-mediated endocytosis”, “negative regulation of lipoprotein lipase activity”, “negative regulation of lipoprotein particle clearance”, and “positive regulation of receptor binding” (Figure 4E). KEGG pathways such as “herpes simplex virus 1 infection”, “cholesterol metabolism”, and “antigen processing and presentation” were overrepresented (Figure 4F), suggesting RGC susceptibility related to some biological mechanism such as activity regulation.

### Horizontal Cells

For HCs, RFECV identified 110 key genes (Figure 5A and Supplementary Table 1). The top 20 weighted features are plotted in Figure 5B, and bootstrap RFE confirmed the genes selection stability (Figure 5C). The ensuing geneLbased classifier achieved 86% accuracy, with AUCs of 0.91 (NON), 0.97 (DM), and 0.96 (DR) (Figure 5D). These metrics underscore the discriminative validity of the selected set. Some biological processes contribute to the pathogenesis of DR by affecting HCs function and survival. In addition to the genes *CCL2*, *GNGT1* and *KDM5D* already described before, in the context of DR, oxidative stress is a significant factor leading to retinal cell damage, and gene *GPX3* plays a protective role by mitigating reactive oxygen species accumulation [35].

**Figure 5:**
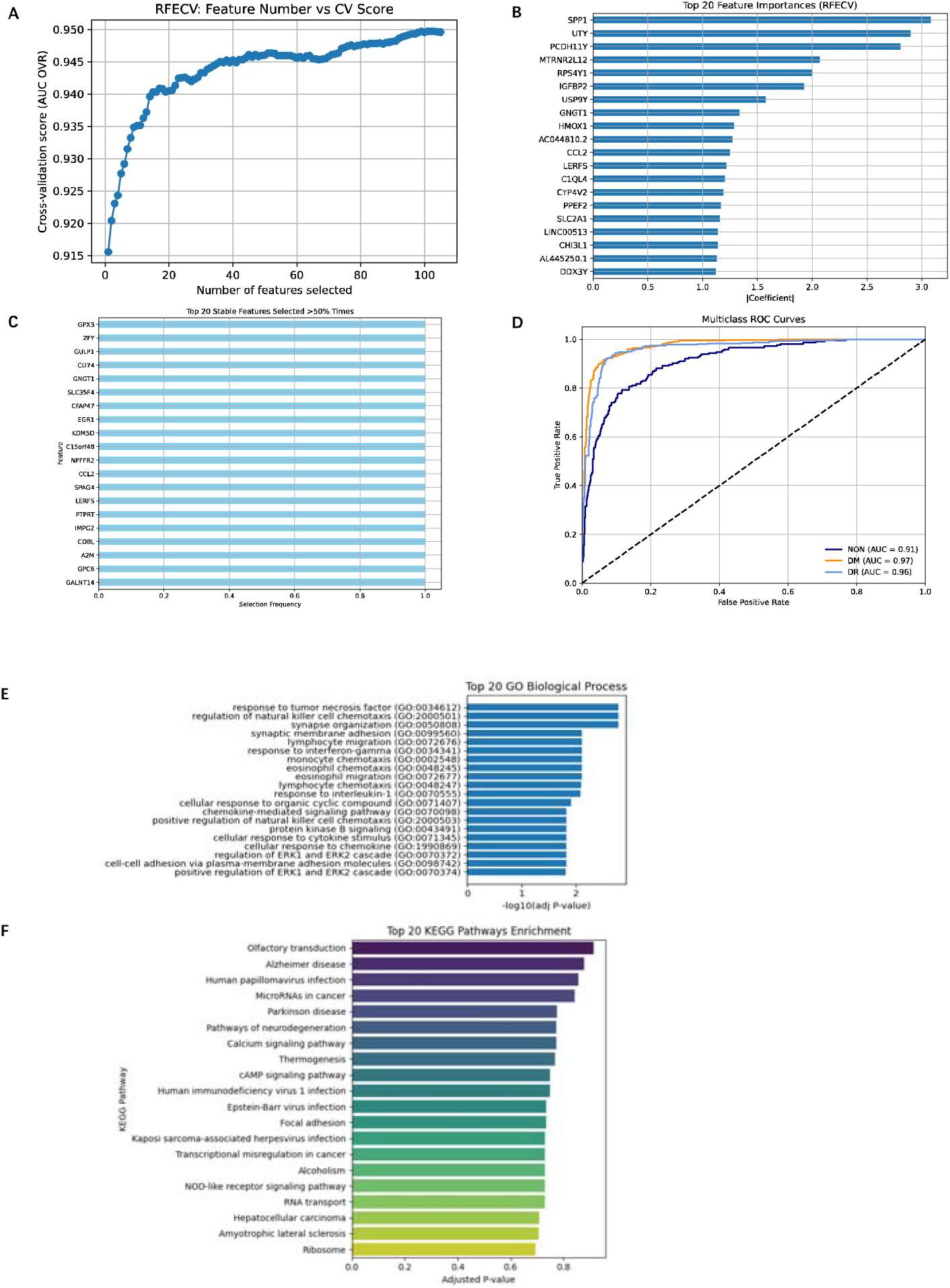
Comprehensive analysis of Chinese human retinal HCs type (A) RFECVLderived feature selection curve: RFECV was performed using an L1-penalized logistic regression classifier under a one-versus-rest scheme. The plot illustrated the relationship between the number of selected features and the corresponding mean cross-validated performance metric (AUC One-vs-Rest). The observed unimodal trend indicated that model performance is maximized with a relatively small feature subset. The optimal feature subset size was therefore chosen for downstream modeling. (B) To assess the relative importance of the features selected by RFECV, we plotted the top 20 features based on the sum of the absolute values of the coefficients across all classes in the logistic regression model. The plot showed the features arranged in descending order of importance. The horizontal bars represent the magnitude of the coefficients, with longer bars indicating greater contribution to the model’s classification performance. Notably, genes such as *SPP1*, *UTY*, and *PCDH11Y* exhibit the highest feature importances, suggesting their critical roles in distinguishing between the different classes (healthy, diabetes, diabetic retinopathy) in this study. (C) To evaluate the robustness of the feature selection process, we performed stability selection by repeating RFE 20 times with different random seeds. Each run selected a fixed number of features, as determined previously by RFECV. For each gene, we calculated the proportion of times it was selected across the 20 repetitions. The plot displayed the top 20 most frequently selected features among those chosen in over 50% of the runs. Higher selection frequency indicates greater stability and reliability of the feature in contributing to classification performance. Genes such as *GPX3*, *ZFY*, and *GULP1* were consistently retained, suggesting their strong and reproducible discriminative power across subsamples. (D) To evaluate the classification performance of the final model, ROC curves were constructed for each class (NON, DM, DR) using a one-versus-rest approach. The plot displayed the true positive rate plotted against the false positive rate for each label. The colored curves represent model performance for each class, with the diagonal dashed line indicating the performance of a random classifier (AUC = 0.5). The area under the curve (AUC) values, reported in the legend, quantify the model’s ability to discriminate each class. All three classes exhibit AUCs substantially greater than 0.5, indicating strong predictive power and reliable differentiation between healthy, diabetic, and retinopathic states. (E) Bar plot presented the top 20 enriched GO terms for the important and stable gene list. Rows represented each GO term that these genes were involved in. The horizontal bars represent the p-value, the longer bars indicating the stronger the correlation. (F) Bar plot presented the top 20 enriched KEGG terms for the important and stable gene list. Rows represented each KEGG term that these genes were involved in. The horizontal bars represent the adjusted p-value, the longer bars indicating the stronger the correlation.

GO analysis revealed enrichment in “response to tumor necrosis factor”, “regulation of natural killer cell chemotaxis”, and “synapse organization” (Figure 5E). KEGG pathways such as “olfactory transduction”, “alzheimer disease”, and “human papillomavirus infection” were also significant (Figure 5F), pointing to some biological process involved like synaptic regulation.

### Cone Photoreceptors

RFECV selection in cone photoreceptors yielded 123 genes (Figure 6A and Supplementary Table 1). Figure 6B presents the top 20 genes by coefficient magnitude, and the genes selected by RFE exhibited high bootstrap stability (Figure 6C). A classifier using these genes reached 91% accuracy, with ROC AUCs of 0.95 (NON), 0.99 (DM), and 0.98 (DR) (Figure 6D). Such performance highlighted the discriminatory capacity of cone photoreceptor key genes. Genes such as *GPX3* and *AK4* are involved in the cellular response to oxidative stress. Gene *AK4* is involved in mitochondrial energy metabolism and may play a role in managing oxidative stress within cone photoreceptors [36]. Genes such as *CCL4L2* and *CD74* are involved in immune and inflammatory pathways [37–38]. Elevated expression of these genes may exacerbate inflammatory responses in the diabetic retina. Genes like *FOSB* and *LERFS* are involved in gene expression regulation, changes in the expression of them can influence cone photoreceptor function under hyperglycemic conditions [39]. Genes such as *TMSB10* and *SLC22A17* are associated with cellular metabolism and structural integrity [40–41].

**Figure 6:**
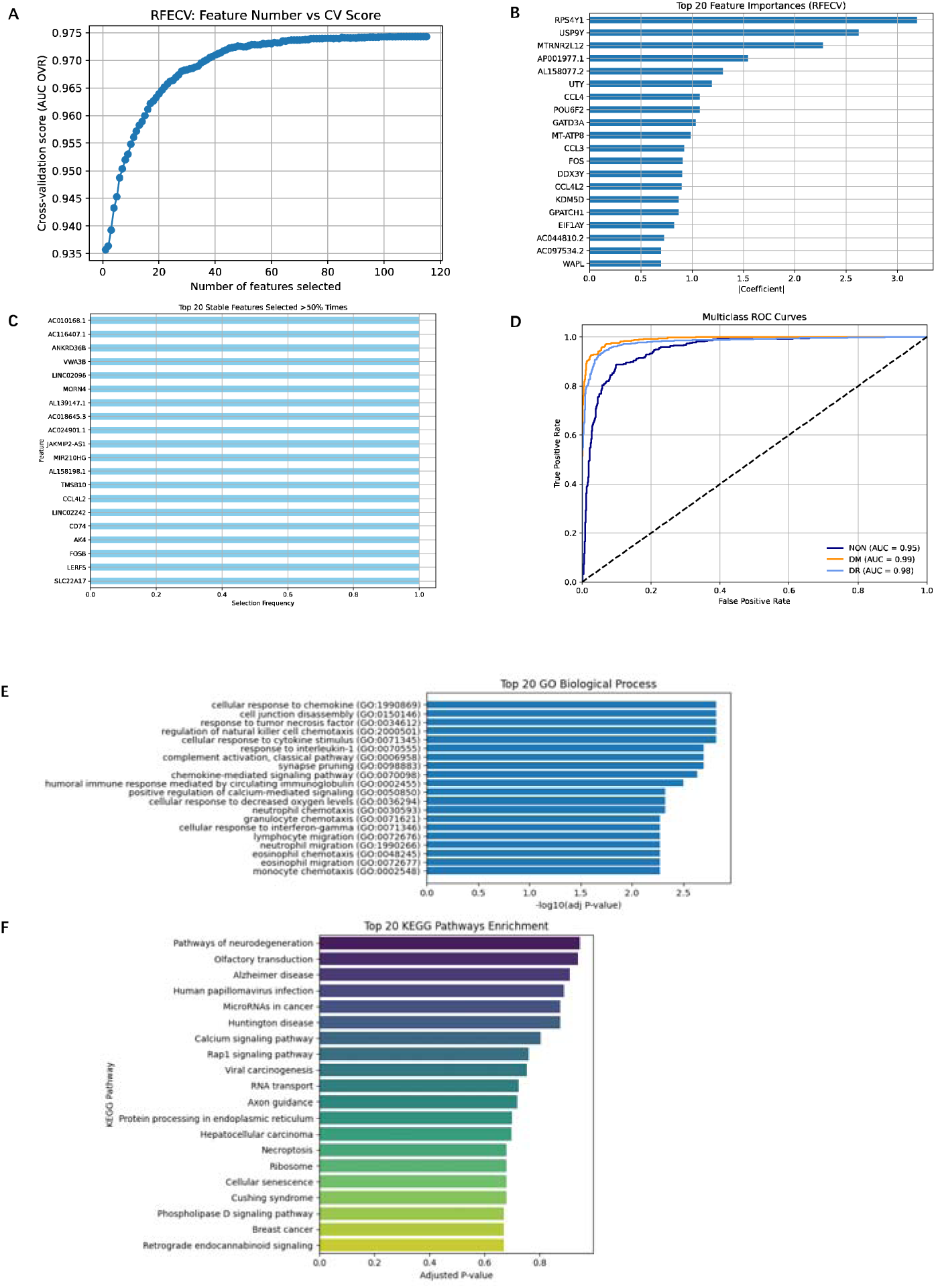
Comprehensive analysis of Chinese human retinal cone photoreceptors type (A) RFECVLderived feature selection curve: RFECV was performed using an L1-penalized logistic regression classifier under a one-versus-rest scheme. The plot illustrated the relationship between the number of selected features and the corresponding mean cross-validated performance metric (AUC One-vs-Rest). The observed unimodal trend indicated that model performance is maximized with a relatively small feature subset. The optimal feature subset size was therefore chosen for downstream modeling. (B) To assess the relative importance of the features selected by RFECV, we plotted the top 20 features based on the sum of the absolute values of the coefficients across all classes in the logistic regression model. The plot showed the features arranged in descending order of importance. The horizontal bars represent the magnitude of the coefficients, with longer bars indicating greater contribution to the model’s classification performance. Notably, genes such as *RPS4Y1*, *USP9Y*, and *MTRNR2L12* exhibit the highest feature importances, suggesting their critical roles in distinguishing between the different classes (healthy, diabetes, diabetic retinopathy) in this study. (C) To evaluate the robustness of the feature selection process, we performed stability selection by repeating RFE 20 times with different random seeds. Each run selected a fixed number of features, as determined previously by RFECV. For each gene, we calculated the proportion of times it was selected across the 20 repetitions. The plot displayed the top 20 most frequently selected features among those chosen in over 50% of the runs. Higher selection frequency indicates greater stability and reliability of the feature in contributing to classification performance. Genes such as *AC010168.1*, *AC116407.1*, and *ANKRD36B* were consistently retained, suggesting their strong and reproducible discriminative power across subsamples. (D) To evaluate the classification performance of the final model, ROC curves were constructed for each class (NON, DM, DR) using a one-versus-rest approach. The plot displayed the true positive rate plotted against the false positive rate for each label. The colored curves represent model performance for each class, with the diagonal dashed line indicating the performance of a random classifier (AUC = 0.5). The area under the curve (AUC) values, reported in the legend, quantify the model’s ability to discriminate each class. All three classes exhibit AUCs substantially greater than 0.5, indicating strong predictive power and reliable differentiation between healthy, diabetic, and retinopathic states. (E) Bar plot presented the top 20 enriched GO terms for the important and stable gene list. Rows represented each GO term that these genes were involved in. The horizontal bars represent the p-value, the longer bars indicating the stronger the correlation. (F) Bar plot presented the top 20 enriched KEGG terms for the important and stable gene list. Rows represented each KEGG term that these genes were involved in. The horizontal bars represent the adjusted p-value, the longer bars indicating the stronger the correlation.

GO enrichment underscored “cellular response to chemokine”, “cell junction disassembly”, and “regulation of natural killer cell chemotaxis” (Figure 6E). KEGG pathways “pathways of neurodegeneration”, “olfactory transduction”, and “alzheimer disease” featured prominently (Figure 6F).

### Rod Photoreceptors

In rod photoreceptors, RFECV pinpointed 117 signature genes (Figure 7A and Supplementary Table 1). The leading 20 features with top coefficient magnitude are displayed in Figure 7B, and the genes with high stability is shown in Figure 7C. The rodLspecific classifier achieved 79% accuracy and multiclass AUCs of 0.85 (NON), 0.95 (DM), and 0.92 (DR) (Figure 7D), underscoring exceptional discriminative performance. Genes such as *CD163*, *TYROBP*, and *C1QC* are involved in immune and inflammatory pathways. Gene *HAMP* encodes hepcidin, a key regulator of iron homeostasis, and dysregulation of iron metabolism can lead to oxidative stress that caused rod photoreceptor apoptosis [42]. The expression of gene *EGR3* can influence rod photoreceptor function under hyperglycemic conditions [43]. Genes such as *SLC44A5* and *USP9Y* are associated with cellular metabolism and structural integrity [44].

**Figure 7:**
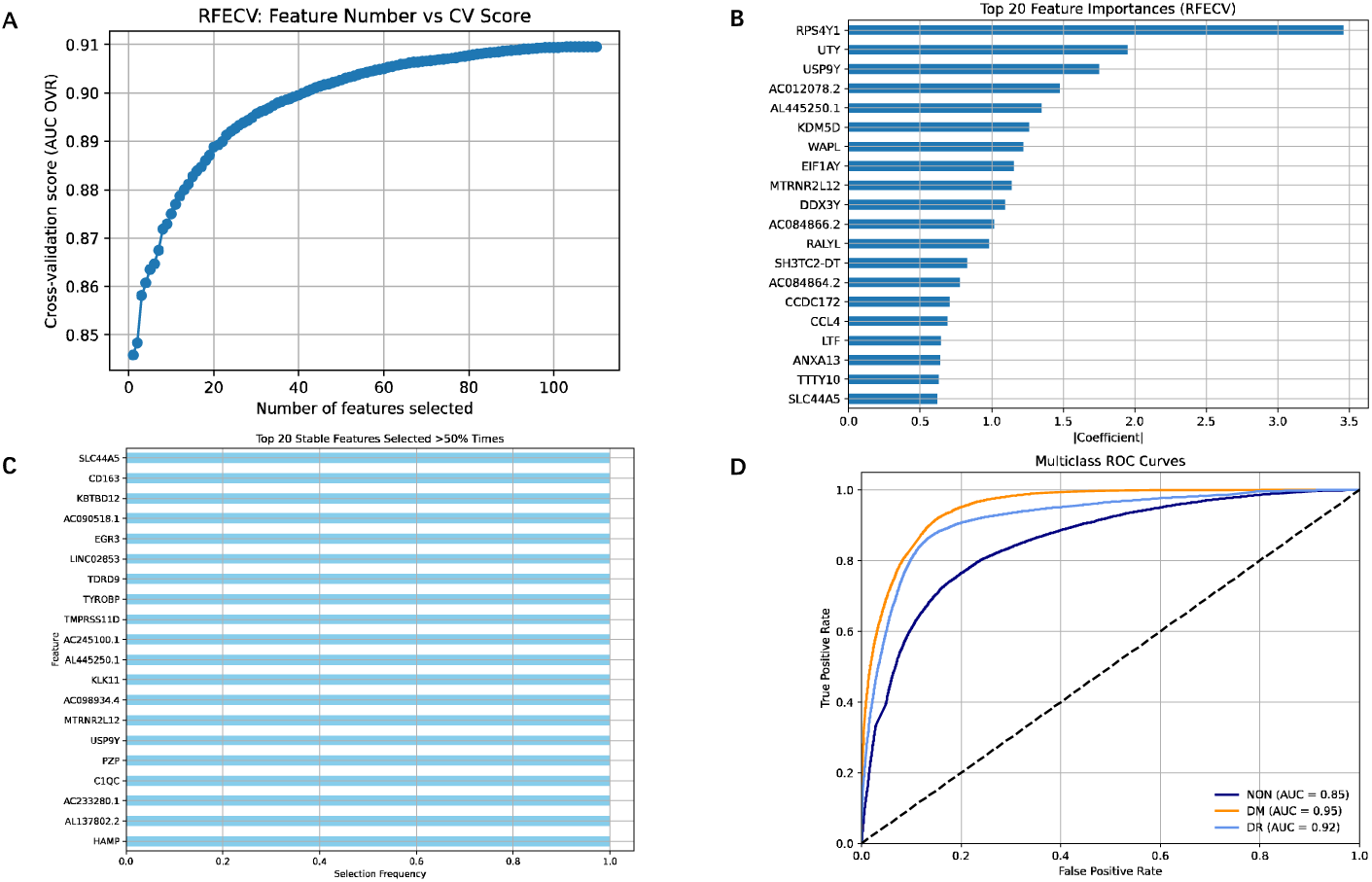
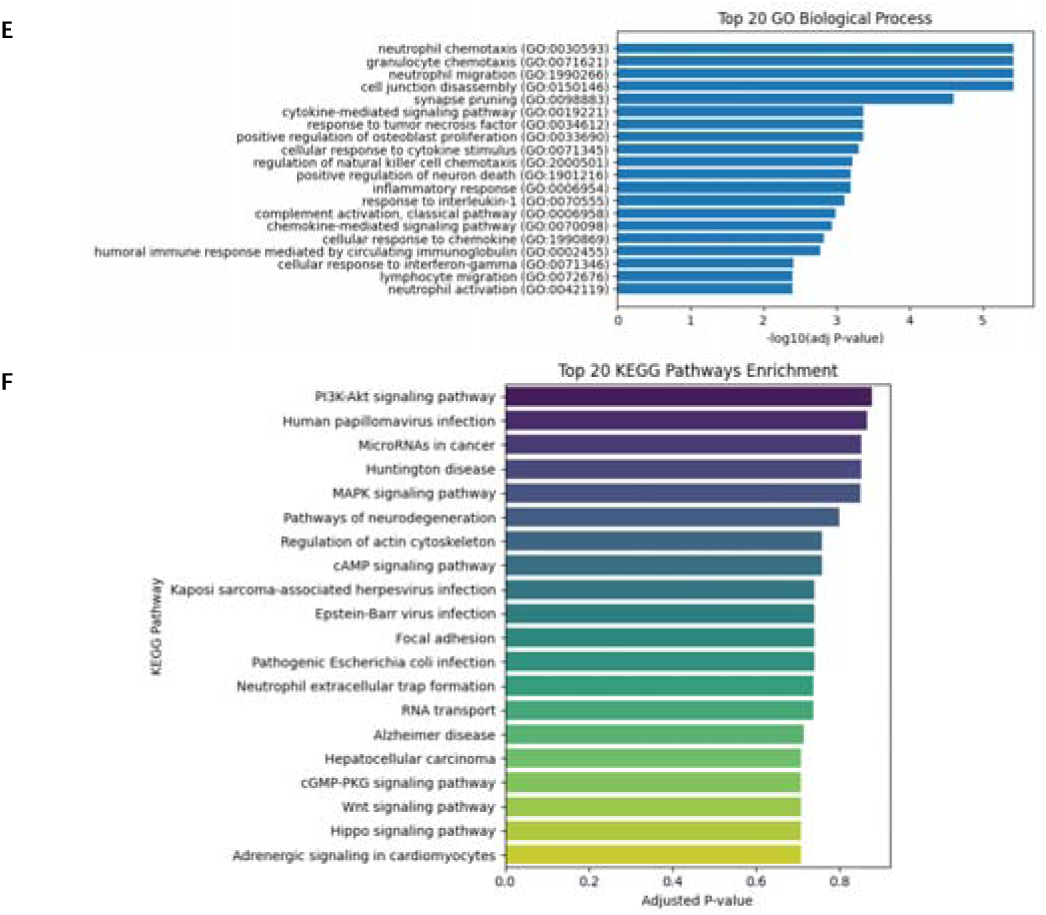
Comprehensive analysis of Chinese human retinal rod photoreceptors (A) RFECVLderived feature selection curve: RFECV was performed using an L1-penalized logistic regression classifier under a one-versus-rest scheme. The plot illustrated the relationship between the number of selected features and the corresponding mean cross-validated performance metric (AUC One-vs-Rest). The observed unimodal trend indicated that model performance is maximized with a relatively small feature subset. The optimal feature subset size was therefore chosen for downstream modeling. (B) To assess the relative importance of the features selected by RFECV, we plotted the top 20 features based on the sum of the absolute values of the coefficients across all classes in the logistic regression model. The plot showed the features arranged in descending order of importance. The horizontal bars represent the magnitude of the coefficients, with longer bars indicating greater contribution to the model’s classification performance. Notably, genes such as *RPS4Y1*, *UTY*, and *USP9Y* exhibit the highest feature importances, suggesting their critical roles in distinguishing between the different classes (healthy, diabetes, diabetic retinopathy) in this study. (C) To evaluate the robustness of the feature selection process, we performed stability selection by repeating RFE 20 times with different random seeds. Each run selected a fixed number of features, as determined previously by RFECV. For each gene, we calculated the proportion of times it was selected across the 20 repetitions. The plot displayed the top 20 most frequently selected features among those chosen in over 50% of the runs. Higher selection frequency indicates greater stability and reliability of the feature in contributing to classification performance. Genes such as *SLC44A5*, *CD163*, and *KBTBD12* were consistently retained, suggesting their strong and reproducible discriminative power across subsamples. (D) To evaluate the classification performance of the final model, ROC curves were constructed for each class (NON, DM, DR) using a one-versus-rest approach. The plot displayed the true positive rate plotted against the false positive rate for each label. The colored curves represent model performance for each class, with the diagonal dashed line indicating the performance of a random classifier (AUC = 0.5). The area under the curve (AUC) values, reported in the legend, quantify the model’s ability to discriminate each class. All three classes exhibit AUCs substantially greater than 0.5, indicating strong predictive power and reliable differentiation between healthy, diabetic, and retinopathic states. (E) Bar plot presented the top 20 enriched GO terms for the important and stable gene list. Rows represented each GO term that these genes were involved in. The horizontal bars represent the p-value, the longer bars indicating the stronger the correlation. (F) Bar plot presented the top 20 enriched KEGG terms for the important and stable gene list. Rows represented each KEGG term that these genes were involved in. The horizontal bars represent the adjusted p-value, the longer bars indicating the stronger the correlation.

GO analysis revealed “neutrophil chemotaxis”, “granulocyte chemotaxis”, and “neutrophil migration” (Figure 7E). KEGG pathways “PI3K-Akt signaling pathway”, “human papillomavirus infection”, and “microRNAs in cancer” were highly enriched (Figure 7F).

### Microglia

For microglia, RFECV selected 93 genes (Figure 8A and Supplementary Table 1). The top 20 features by weight are shown in Figure 8B, with bootstrap validation in Figure 8C confirming their consistency. A classifier trained on these genes yielded 84% accuracy and AUCs of 0.93 (NON), 0.96 (DM), and 0.93 (DR) (Figure 8D), reflecting moderate discriminative power. Genes such as *IL6* and *IGFBP5* are involved in mediating inflammation within the diabetic retina [45]. Genes like *HMOX* and *MT1E* are associated with the cellular response to oxidative stress [46–47]. Genes *FN1* and *SDC3* are involved in extracellular matrix organization [48–49]. Genes *FGF13* and *PTPN13* are implicated in neurovascular functions [50–51].

**Figure 8:**
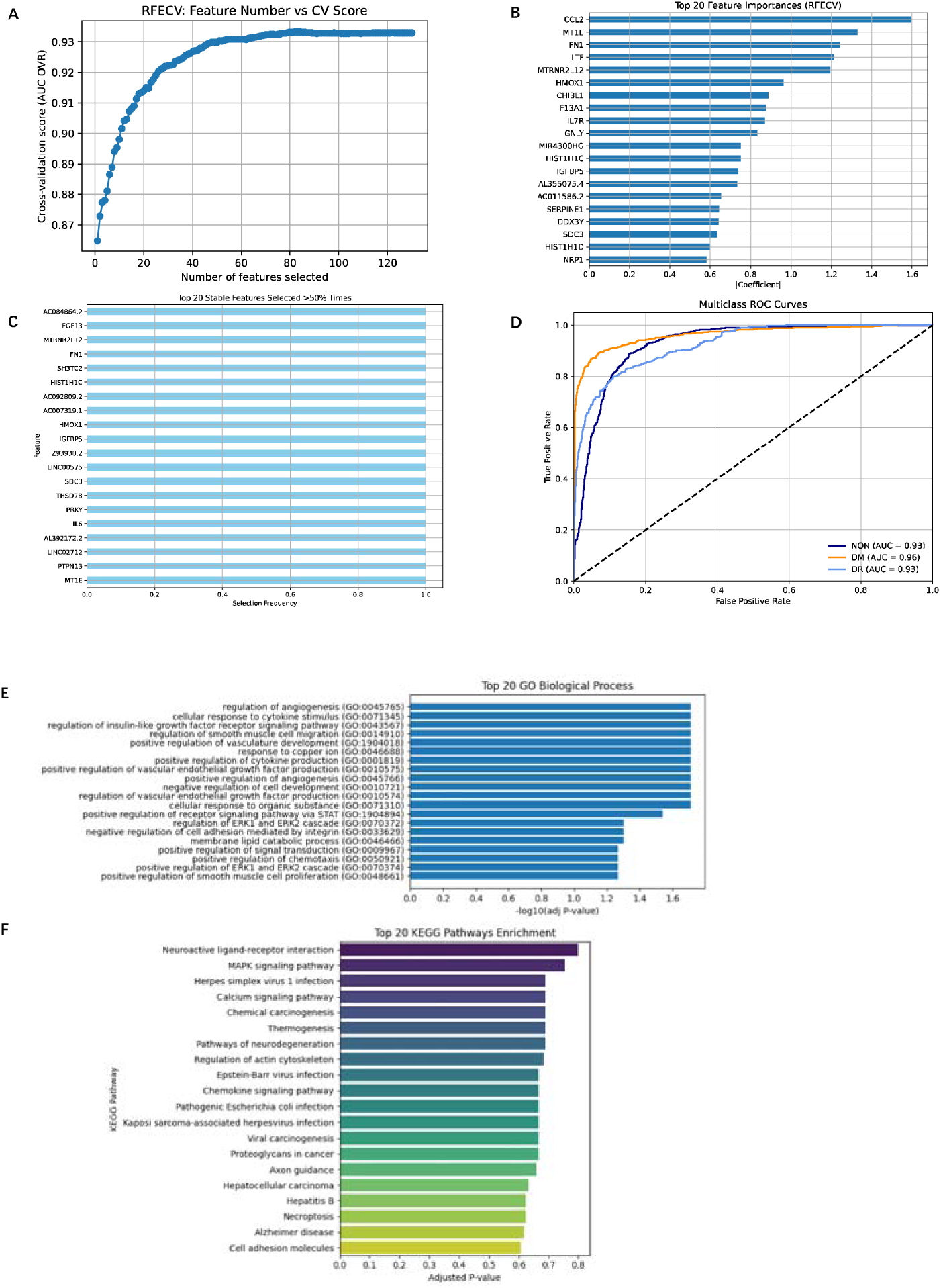
Comprehensive analysis of Chinese human retinal microglia type (A) RFECVLderived feature selection curve: RFECV was performed using an L1-penalized logistic regression classifier under a one-versus-rest scheme. The plot illustrated the relationship between the number of selected features and the corresponding mean cross-validated performance metric (AUC One-vs-Rest). The observed unimodal trend indicated that model performance is maximized with a relatively small feature subset. The optimal feature subset size was therefore chosen for downstream modeling. (B) To assess the relative importance of the features selected by RFECV, we plotted the top 20 features based on the sum of the absolute values of the coefficients across all classes in the logistic regression model. The plot showed the features arranged in descending order of importance. The horizontal bars represent the magnitude of the coefficients, with longer bars indicating greater contribution to the model’s classification performance. Notably, genes such as *CCL2*, *MT1E*, and *FN1* exhibit the highest feature importances, suggesting their critical roles in distinguishing between the different classes (healthy, diabetes, diabetic retinopathy) in this study. (C) To evaluate the robustness of the feature selection process, we performed stability selection by repeating RFE 20 times with different random seeds. Each run selected a fixed number of features, as determined previously by RFECV. For each gene, we calculated the proportion of times it was selected across the 20 repetitions. The plot displayed the top 20 most frequently selected features among those chosen in over 50% of the runs. Higher selection frequency indicates greater stability and reliability of the feature in contributing to classification performance. Genes such as *AC084864.2*, *FGF13*, and *MTRNR2L12* were consistently retained, suggesting their strong and reproducible discriminative power across subsamples. (D) To evaluate the classification performance of the final model, ROC curves were constructed for each class (NON, DM, DR) using a one-versus-rest approach. The plot displayed the true positive rate plotted against the false positive rate for each label. The colored curves represent model performance for each class, with the diagonal dashed line indicating the performance of a random classifier (AUC = 0.5). The area under the curve (AUC) values, reported in the legend, quantify the model’s ability to discriminate each class. All three classes exhibit AUCs substantially greater than 0.5, indicating strong predictive power and reliable differentiation between healthy, diabetic, and retinopathic states. (E) Bar plot presented the top 20 enriched GO terms for the important and stable gene list. Rows represented each GO term that these genes were involved in. The horizontal bars represent the p-value, the longer bars indicating the stronger the correlation. (F) Bar plot presented the top 20 enriched KEGG terms for the important and stable gene list. Rows represented each KEGG term that these genes were involved in. The horizontal bars represent the adjusted p-value, the longer bars indicating the stronger the correlation.

GO enrichment emphasized “regulation of angiogenesis”, “cellular response to cytokine stimulus”, and “regulation of insulin-like growth factor receptor signaling pathway” (Figure 8E). KEGG pathways including “neuroactive ligand-receptor interaction”, “MAPK signaling pathway”, and “calcium signaling pathway” were enriched (Figure 8F).

### Astrocytes

In astrocytes, RFECV isolated 61 key genes (Figure 9A and Supplementary Table 1). The top 20 by coefficient magnitude are plotted in Figure 9B, and the genes with high bootstrap repeatability are shown in Figure 9C. The astrocyteLfocused classifier achieved 81% accuracy and AUCs of 0.89 (NON), 0.96 (DM), and 0.95 (DR) (Figure 9D). Genes such as *SPP1* and *CCL3* are involved in mediating inflammation within the diabetic retina [52–53]. Genes like *MT-ATP8* and *CRABP1* are associated with the cellular response to oxidative stress [54–55]. Gene *PER1* is involved in the regulation of circadian rhythms and altered expression of it suggests that diabetes disrupts the normal circadian regulation in retinal cells [56]. Genes such as *KLF4* and *SOX2* are transcription factors that regulate astrocyte differentiation and function [57–58].

**Figure 9:**
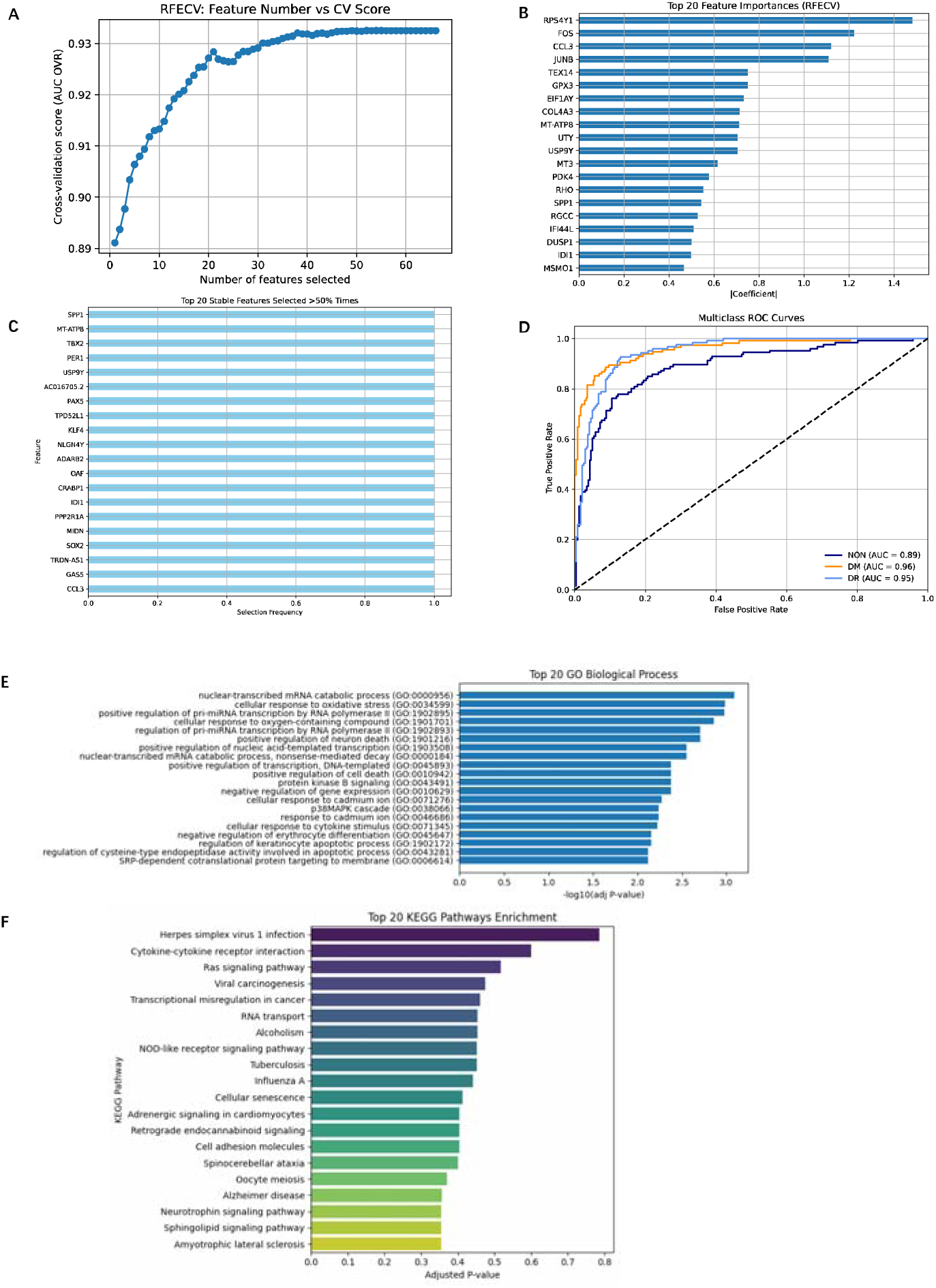
Comprehensive analysis of Chinese human retinal astrocyte type (A) RFECVLderived feature selection curve: RFECV was performed using an L1-penalized logistic regression classifier under a one-versus-rest scheme. The plot illustrated the relationship between the number of selected features and the corresponding mean cross-validated performance metric (AUC One-vs-Rest). The observed unimodal trend indicated that model performance is maximized with a relatively small feature subset. The optimal feature subset size was therefore chosen for downstream modeling. (B) To assess the relative importance of the features selected by RFECV, we plotted the top 20 features based on the sum of the absolute values of the coefficients across all classes in the logistic regression model. The plot showed the features arranged in descending order of importance. The horizontal bars represent the magnitude of the coefficients, with longer bars indicating greater contribution to the model’s classification performance. Notably, genes such as *RPS4Y1*, *FOS*, and *CCL3* exhibit the highest feature importances, suggesting their critical roles in distinguishing between the different classes (healthy, diabetes, diabetic retinopathy) in this study. (C) To evaluate the robustness of the feature selection process, we performed stability selection by repeating RFE 20 times with different random seeds. Each run selected a fixed number of features, as determined previously by RFECV. For each gene, we calculated the proportion of times it was selected across the 20 repetitions. The plot displayed the top 20 most frequently selected features among those chosen in over 50% of the runs. Higher selection frequency indicates greater stability and reliability of the feature in contributing to classification performance. Genes such as *SPP1*, *MT-ATP8*, and *TBX2* were consistently retained, suggesting their strong and reproducible discriminative power across subsamples. (D) To evaluate the classification performance of the final model, ROC curves were constructed for each class (NON, DM, DR) using a one-versus-rest approach. The plot displayed the true positive rate plotted against the false positive rate for each label. The colored curves represent model performance for each class, with the diagonal dashed line indicating the performance of a random classifier (AUC = 0.5). The area under the curve (AUC) values, reported in the legend, quantify the model’s ability to discriminate each class. All three classes exhibit AUCs substantially greater than 0.5, indicating strong predictive power and reliable differentiation between healthy, diabetic, and retinopathic states. (E) Bar plot presented the top 20 enriched GO terms for the important and stable gene list. Rows represented each GO term that these genes were involved in. The horizontal bars represent the p-value, the longer bars indicating the stronger the correlation. (F) Bar plot presented the top 20 enriched KEGG terms for the important and stable gene list. Rows represented each KEGG term that these genes were involved in. The horizontal bars represent the adjusted p-value, the longer bars indicating the stronger the correlation.

GO terms enriched in these genes include “nuclear-transcribed mRNA catabolic process”, “cellular response to oxidative stress”, and “positive regulation of pre-miRNA transcription by RNA polymerase II” (Figure 9E). KEGG pathways such as “herpes simplex virus 1 infection”, “cytokine-cytokine receptor interaction” and “Ras signaling pathway” were significant (Figure 9F), underscoring astrocytic roles in redox homeostasis and metabolic adaptation.

### Müller Glial Cells

RFECV selection in MGCs identified 127 genes (Figure 10A and Supplementary Table 1). Figure 10B depicts the top 20 features, all of which exhibited high bootstrapped stability (Figure 10C). The corresponding classifier achieved 81% accuracy and AUCs of 0.88 (NON), 0.97 (DM), and 0.95 (DR) (Figure 10D), highlighting exceptional classification performance. Genes such as *GIMAP7* and *KISS1R* are involved in immune regulation and inflammatory pathways [59–60]. Gene *GSTM5* and *CYP26A1* are associated with oxidative stress and metabolic regulation [61–62]. Several long non-coding RNAs such as *AL583808.1*, *ZBTB20-AS2*, and *AC018628.2* are involved in gene regulation including chromatin remodeling, transcription, and post-transcriptional processing, which influences MGCs function under diabetic conditions [63].

**Figure 10:**
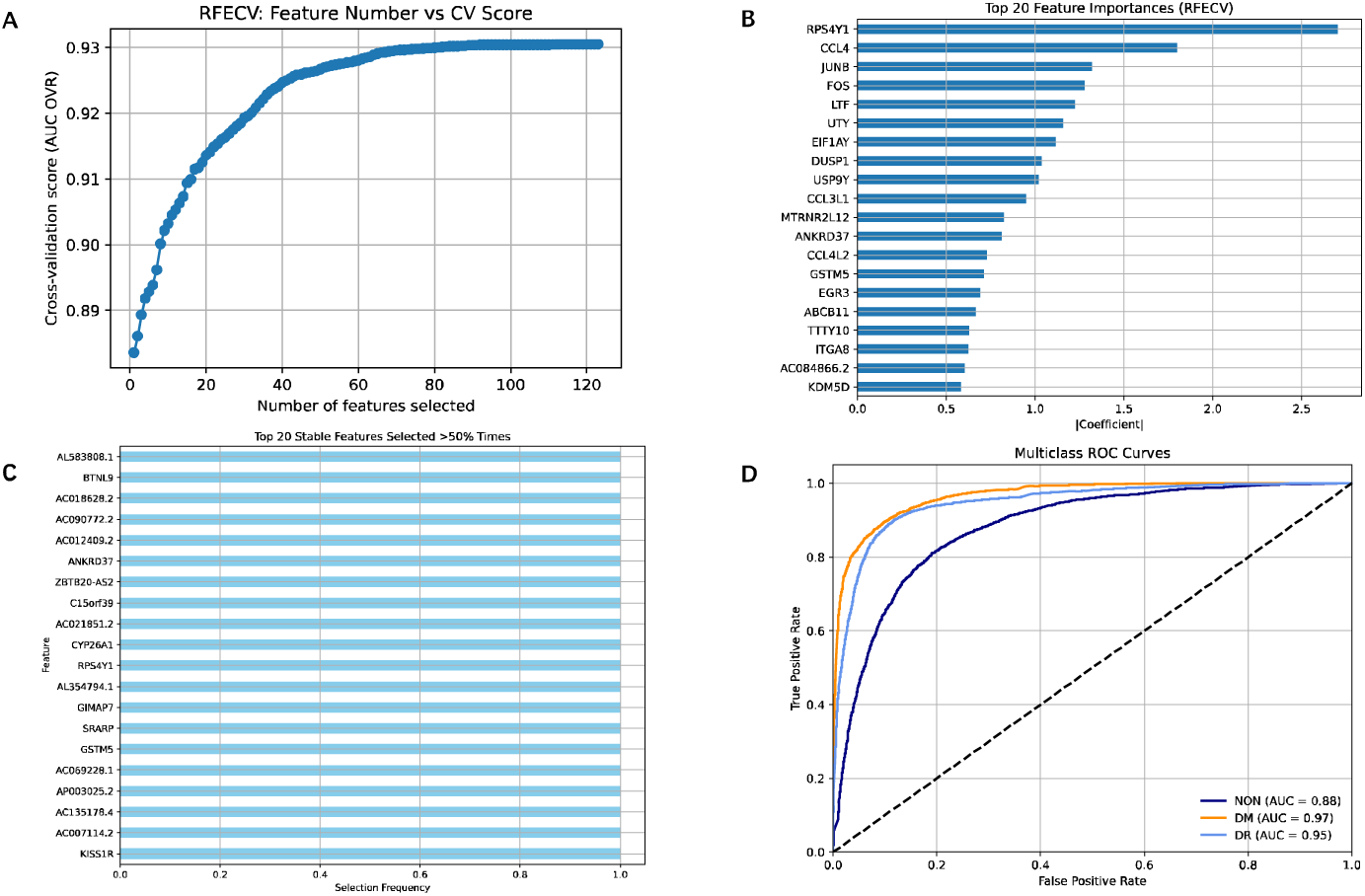
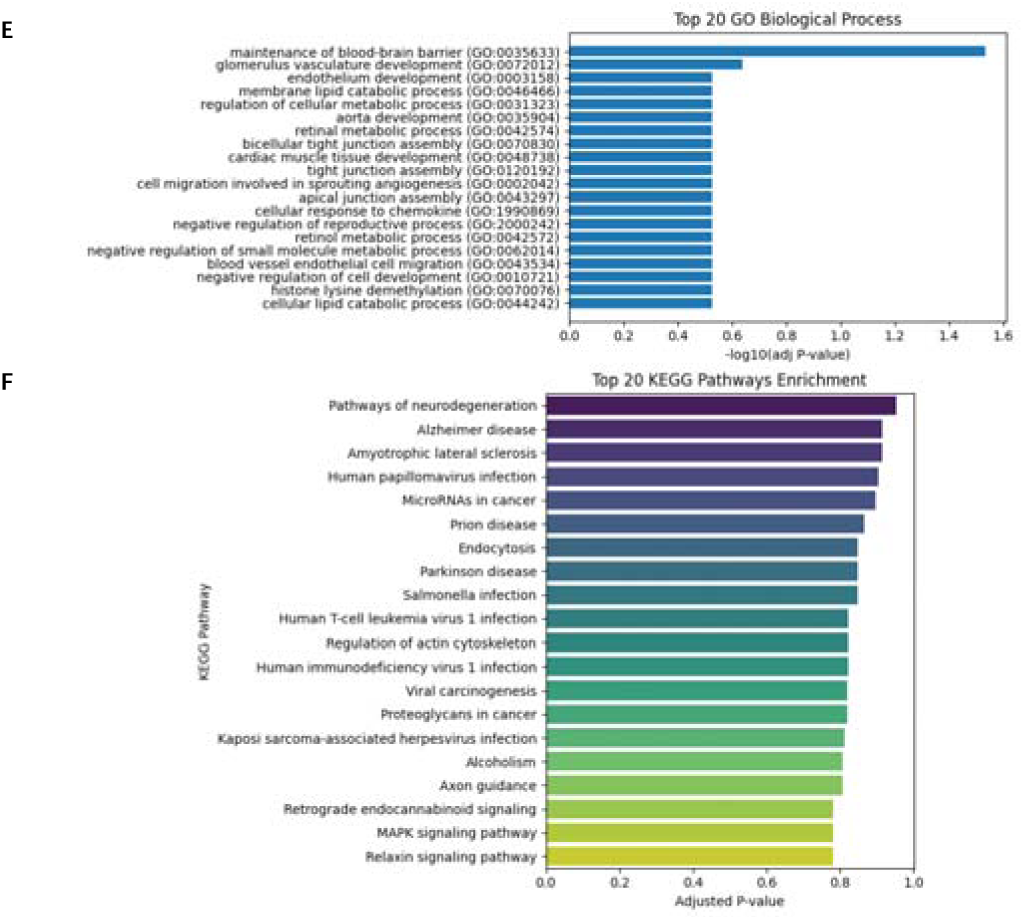
Comprehensive analysis of Chinese human retinal MGCs type (A) RFECVLderived feature selection curve: RFECV was performed using an L1-penalized logistic regression classifier under a one-versus-rest scheme. The plot illustrated the relationship between the number of selected features and the corresponding mean cross-validated performance metric (AUC One-vs-Rest). The observed unimodal trend indicated that model performance is maximized with a relatively small feature subset. The optimal feature subset size was therefore chosen for downstream modeling. (B) To assess the relative importance of the features selected by RFECV, we plotted the top 20 features based on the sum of the absolute values of the coefficients across all classes in the logistic regression model. The plot showed the features arranged in descending order of importance. The horizontal bars represent the magnitude of the coefficients, with longer bars indicating greater contribution to the model’s classification performance. Notably, genes such as *RPS4Y1*, *CCL4*, and *JUNB* exhibit the highest feature importances, suggesting their critical roles in distinguishing between the different classes (healthy, diabetes, diabetic retinopathy) in this study. (C) To evaluate the robustness of the feature selection process, we performed stability selection by repeating RFE 20 times with different random seeds. Each run selected a fixed number of features, as determined previously by RFECV. For each gene, we calculated the proportion of times it was selected across the 20 repetitions. The plot displayed the top 20 most frequently selected features among those chosen in over 50% of the runs. Higher selection frequency indicates greater stability and reliability of the feature in contributing to classification performance. Genes such as *AL583808.1*, *BTNL9*, and *AC018628.2* were consistently retained, suggesting their strong and reproducible discriminative power across subsamples. (D) To evaluate the classification performance of the final model, ROC curves were constructed for each class (NON, DM, DR) using a one-versus-rest approach. The plot displayed the true positive rate plotted against the false positive rate for each label. The colored curves represent model performance for each class, with the diagonal dashed line indicating the performance of a random classifier (AUC = 0.5). The area under the curve (AUC) values, reported in the legend, quantify the model’s ability to discriminate each class. All three classes exhibit AUCs substantially greater than 0.5, indicating strong predictive power and reliable differentiation between healthy, diabetic, and retinopathic states. (E) Bar plot presented the top 20 enriched GO terms for the important and stable gene list. Rows represented each GO term that these genes were involved in. The horizontal bars represent the p-value, the longer bars indicating the stronger the correlation. (F) Bar plot presented the top 20 enriched KEGG terms for the important and stable gene list. Rows represented each KEGG term that these genes were involved in. The horizontal bars represent the adjusted p-value, the longer bars indicating the stronger the correlation.

GO enrichment revealed “maintenance of blood-brain barrier”, “glomerulus vasculature development”, and “endothelium development” (Figure 10E). KEGG pathways including “pathways of neurodegeneration”, “alzheimer disease”, and “amyotrophic lateral sclerosis” were enriched (Figure 10F), implicating MGCs in structural remodeling and gliosis.

### T Cells

Finally, in retinal-infiltrating T cells, RFECV selected 44 discriminatory genes (Figure 11A; Supplementary Table 1). The leading 20 features are displayed in Figure 11B, with bootstrap stability confirmed in Figure 11C. The T Cell classifier achieved 95% accuracy, with AUCs of 0.97 (NON), 0.99 (DM), and 0.98 (DR) (Figure 11D). The identified important genes in T cells are implicated in various biological processes. Genes such as *CD83*, *HLA-DRA*, and *BCL3* are involved in immune regulation and inflammatory pathways [64–66]. Genes like *MT2A*, *MT1X*, and *MT-ND4L* are associated with mitochondrial function and oxidative stress responses [47]. Genes such as *NRL*, *CRX*, *RHO*, *SAG*, and *RTBDN* are traditionally associated with photoreceptor development and function [67–68]. The presence of these genes in T cells may indicate aberrant gene expression or potential interactions between immune cells and photoreceptor pathways in DR.

**Figure 11:**
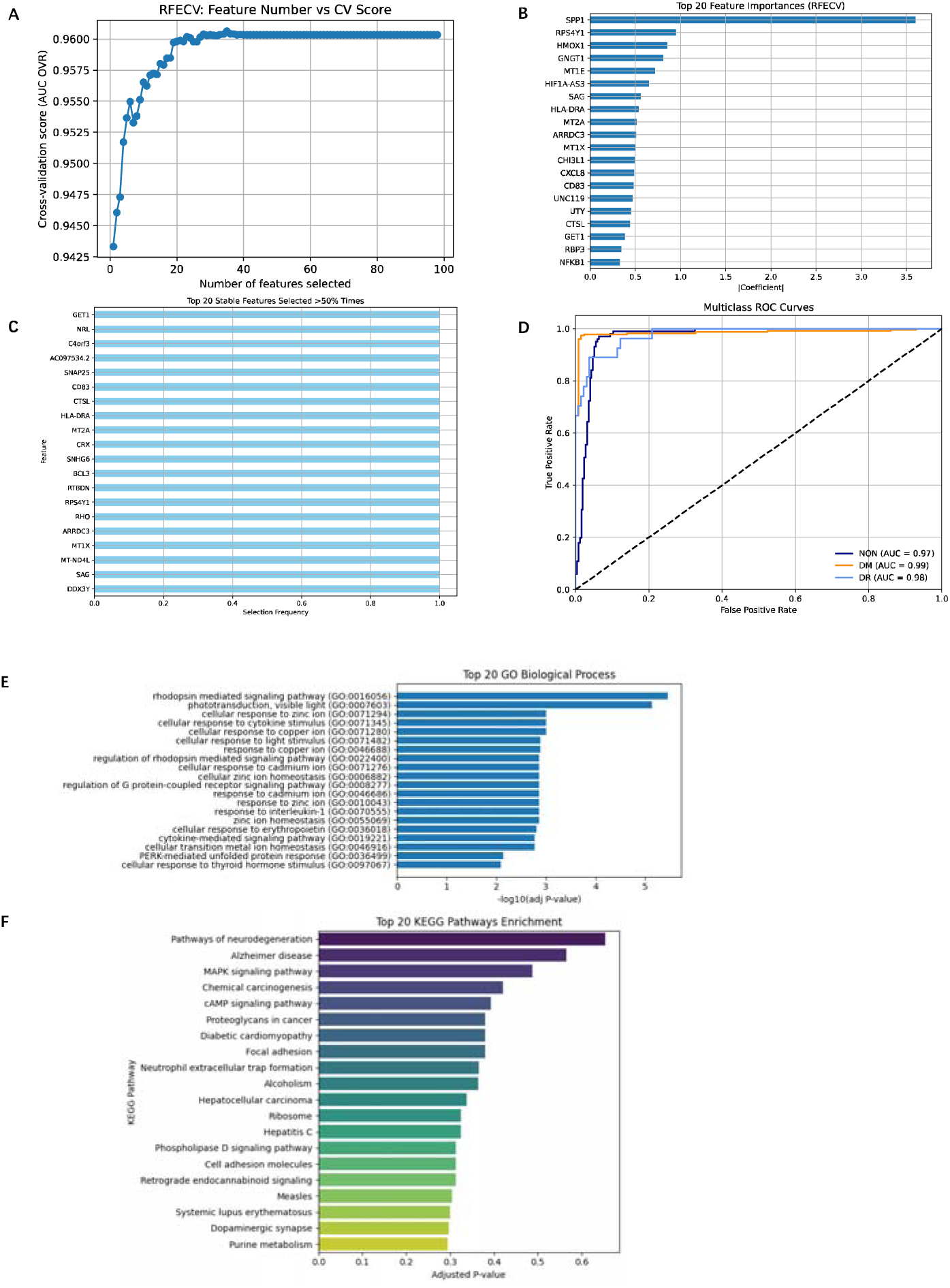
Comprehensive analysis of Chinese human retinal T cells type (A) RFECVLderived feature selection curve: RFECV was performed using an L1-penalized logistic regression classifier under a one-versus-rest scheme. The plot illustrated the relationship between the number of selected features and the corresponding mean cross-validated performance metric (AUC One-vs-Rest). The observed unimodal trend indicated that model performance is maximized with a relatively small feature subset. The optimal feature subset size was therefore chosen for downstream modeling. (B) To assess the relative importance of the features selected by RFECV, we plotted the top 20 features based on the sum of the absolute values of the coefficients across all classes in the logistic regression model. The plot showed the features arranged in descending order of importance. The horizontal bars represent the magnitude of the coefficients, with longer bars indicating greater contribution to the model’s classification performance. Notably, genes such as *SSP1*, *RPS4Y1*, and *HMOX1* exhibit the highest feature importances, suggesting their critical roles in distinguishing between the different classes (healthy, diabetes, diabetic retinopathy) in this study. (C) To evaluate the robustness of the feature selection process, we performed stability selection by repeating RFE 20 times with different random seeds. Each run selected a fixed number of features, as determined previously by RFECV. For each gene, we calculated the proportion of times it was selected across the 20 repetitions. The plot displayed the top 20 most frequently selected features among those chosen in over 50% of the runs. Higher selection frequency indicates greater stability and reliability of the feature in contributing to classification performance. Genes such as *GET1*, *NRL*, and *C4orf3* were consistently retained, suggesting their strong and reproducible discriminative power across subsamples. (D) To evaluate the classification performance of the final model, ROC curves were constructed for each class (NON, DM, DR) using a one-versus-rest approach. The plot displayed the true positive rate plotted against the false positive rate for each label. The colored curves represent model performance for each class, with the diagonal dashed line indicating the performance of a random classifier (AUC = 0.5). The area under the curve (AUC) values, reported in the legend, quantify the model’s ability to discriminate each class. All three classes exhibit AUCs substantially greater than 0.5, indicating strong predictive power and reliable differentiation between healthy, diabetic, and retinopathic states. (E) Bar plot presented the top 20 enriched GO terms for the important and stable gene list. Rows represented each GO term that these genes were involved in. The horizontal bars represent the p-value, the longer bars indicating the stronger the correlation. (F) Bar plot presented the top 20 enriched KEGG terms for the important and stable gene list. Rows represented each KEGG term that these genes were involved in. The horizontal bars represent the adjusted p-value, the longer bars indicating the stronger the correlation.

GO terms significantly enriched comprised “rhodopsin mediated signaling pathway”, “phototransduction, visible light”, and “cellular response to zinc ion” (Figure 11E). KEGG pathways such as “pathways of neurodegeneration”, “alzheimer disease”, and “MAPK signaling pathway” were overrepresented (Figure 11F).

### Neural Network-Based General Classification Model

We derived ten lists of important genes—one from each of ten distinct retinal cell types—and subsequently merged them to form a consolidated gene list. After removing redundancies, this aggregate list comprised 567 unique genes (Supplementary Table 1). Leveraging these 567 gene signatures, we trained a general neural network classification model capable of classifying and predicting disease states in human retinal cells irrespective of their specific cell-type identity. Within the consolidated gene set, we observed the following recurrence patterns across the ten cell-type–specific important gene lists: two genes (*USP9Y* and *DDX3Y*) were shared by all ten cell types; two genes (*RPS4Y1* and *UTY*) appeared in nine lists; one gene (*ZFY*) appeared in eight; seven genes recurred in seven lists; another seven in six lists; nine in five lists; eleven in four lists; nineteen in three lists; sixty in two lists; and the remainder were unique to a single cell-type list.

We developed and evaluated a general-purpose feed-forward neural network classifier to distinguish among three retinal health states (NON, DM, and DR), using single-cell high-dimensional transcriptomic data. The expression matrix for the consolidated gene list of important genes was extracted, and the expression matrix data was partitioned into stratified training (70%) and test (30%) subsets. To ensure that all gene features contributed comparably to gradient-based optimization, we applied standard scaling to zero-center and unit-variance normalize each gene feature across the data set before transforming both training and test matrices. We then developed a multilayer perceptron (MLP) with two hidden layers of 100 neurons each, using ReLU activations, an adaptive learning-rate Adam optimizer. Training proceeded for up to 200 iterations, with early convergence naturally accommodated by the learning-rate adaptation. On the held-out test set (NL=L89,137 cells), the classifier achieved an overall accuracy of 95.31% and class-specific performance metrics were showed in the Figure 12. Macro-averaged and weighted-averaged FL-scores both reached 0.95, indicating balanced performance across classes despite differing class prevalences. These results demonstrated that the proposed MLP architecture can robustly discriminate among healthy and disease conditions in single-cell transcriptomic data, providing a scalable framework for downstream biomarker discovery and cellLstate mapping in diabetic retinal diseases.

**Figure 12:**
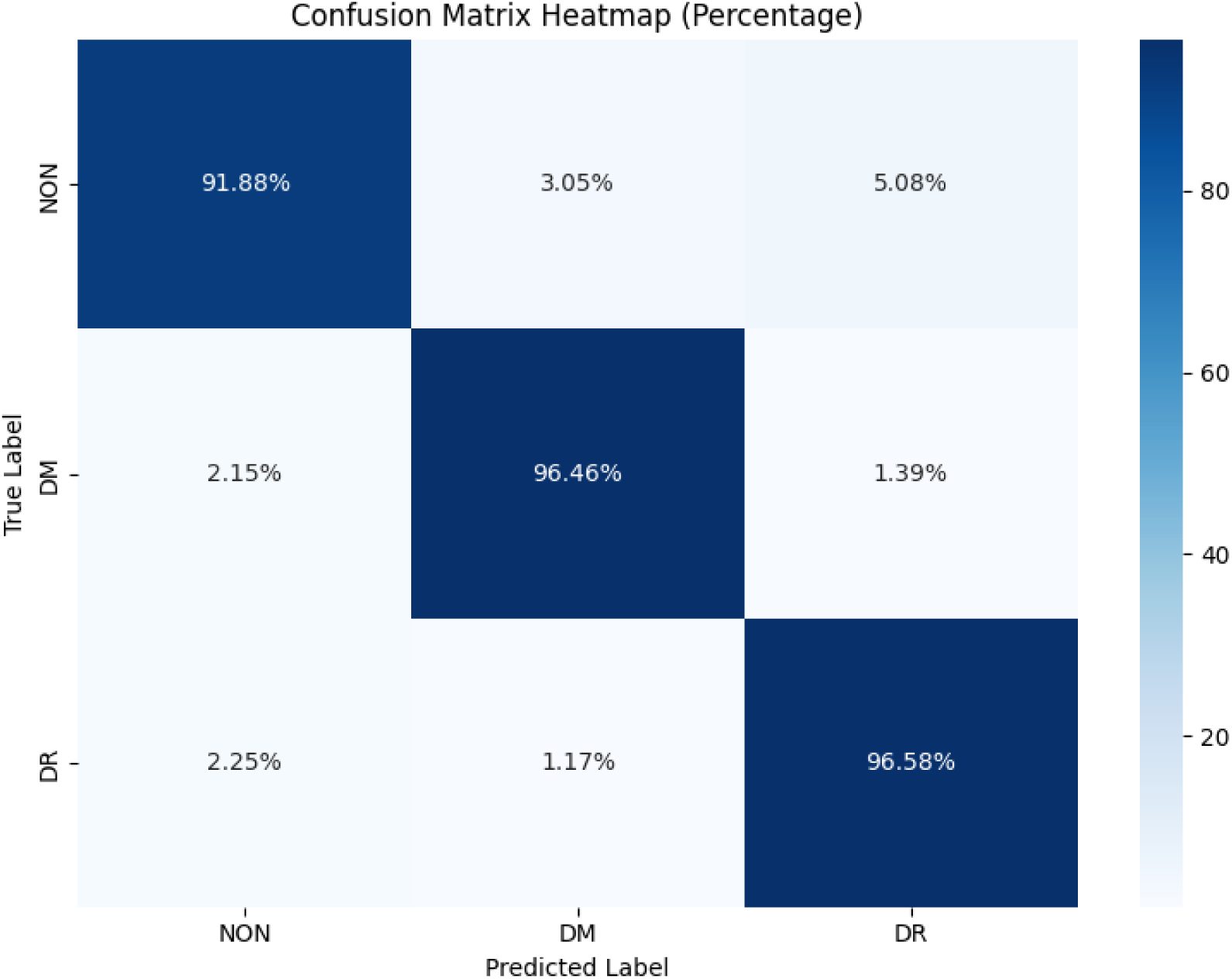
Confusion matrix heatmap of MLP To evaluate the classification accuracy, a confusion matrix heatmap was constructed, where the rows correspond to the actual class labels and the columns to the predicted labels. Each cell contains the percentage of instances for a given actual-predicted class pair, with a color gradient indicating the frequency. 91.88% of NON (samples without diabetes) cells, 96.46% of DM (samples with diabetes) cells and 96.58% of DR (samples with diabetic retinopathy) cells were predicted to the accurate class. The heatmap reveals strong diagonal dominance, indicating high overall classification accuracy (95.31%), while off-diagonal entries highlight specific misclassifications between closely related classes.

## Discussion

In this study, we constructed a high-resolution single-cell transcriptomic atlas of the Chinese human retina, integrating data from donors with diabetic, diabetic retinopathy and non-diabetic to elucidate the cellular and molecular underpinnings of DR. Our integration of state-of-the-art high-resolution scRNA-seq profiling with rigorous machine learning–based feature selection and classification strategies, we systematically identified cell-type–specific genes that discriminate diabetic disease states with high accuracy and biological interpretability, and uncovered disease-relevant pathways.

Our results revealed profound transcriptional heterogeneity across major retinal cell types in response to DR. Notably, inflammation, oxidative stress, immune activation, and neurodegeneration-associated pathways were consistently enriched across diverse cell populations, including ACs, BCs, photoreceptors, and glial cells. The convergence of these pathological signatures in distinct cell types underscores the multifaceted nature of DR and its complex interplay between neurovascular and immune dysfunction. For example, across multiple retinal cell types, we observed consistent upregulation of genes involved in inflammatory pathways, oxidative stress response, and synaptic dysfunction. Recurrent identification of genes such as *CHI3L1*, *CCL2*, *S100A8*, and *GPX3* points to shared cellular responses involving chemokine signaling, immune infiltration, and redox imbalance—hallmarks of chronic diabetic tissue damage. These findings are consistent with previous reports linking chronic low-grade inflammation and oxidative damage to retinal neurovascular degeneration in diabetic patients [69]. Furthermore, the presence of neurodegeneration-related genes and pathways (e.g., KEGG enrichment of “pathways of neurodegeneration” and “MAPK signaling pathway”) across multiple cell types suggests that DR may share mechanistic overlap with other neurodegenerative disorders, such as Alzheimer’s disease and amyotrophic lateral sclerosis. The identification of distinct transcriptional alterations in key retinal cell types—including ACs, BCs, photoreceptors, and glial populations—demonstrates that DR pathophysiology manifests through highly specific cellular mechanisms rather than uniform retinal responses. This observation reinforces the utility of single-cell transcriptomic technique in understanding the intricate cellular perturbations caused by chronic hyperglycemia and inflammation. Furthermore, the recurrence of certain genes (e.g., *DDX3Y*, *USP9Y*, *RPS4Y1*) across multiple retinal cell types suggests the existence of core molecular nodes involved in pan-cellular stress responses or retinal-wide regulatory networks. While the enrichment of Y-linked genes may also reflect underlying sex-based differences in disease susceptibility or expression patterns, this observation merits further investigation in sex-balanced and independent cohorts. One particularly novel finding is the unexpected expression of photoreceptor-related genes (e.g., *RHO*, *SAG*, *CRX*) in infiltrating T cells, raising possibilities regarding cell-cell communication, phagocytic uptake, or ectopic gene expression under diabetic stress. Such results underscore the power of single-cell approaches to uncover noncanonical transcriptional phenomena that may be overlooked in bulk analyses. However, these findings must be interpreted cautiously, and future studies involving spatial transcriptomics or lineage-tracing experiments will be crucial to validate and elucidate these intercellular interactions.

The development of a unified multilayer perceptron (MLP) model trained on an integrated gene set further demonstrated the translational potential of these key genes. Importantly, our application of L1-regularized logistic regression combined with recursive feature elimination and stability selection enabled the extraction of robust and interpretable gene sets with high discriminatory power between DR, DM, and NON states. The subsequent artificial neural network-based classifier trained on the union of selected gene features. Achieving an overall classification accuracy of 95.31%, this model validated the predictive utility of these molecular signatures and exemplified how biologically informed machine learning pipelines can be harnessed to translate high-dimensional transcriptomic data into actionable diagnostic tools. Machine learning–driven analytical frameworks have fundamentally transformed our capacity to interrogate high-dimensional single-cell transcriptomic datasets, enabling the extraction of biologically meaningful patterns that would otherwise remain obscured by noise and technical variability. This framework not only confirms the feasibility of disease-state classification using scRNA-seq data but also provides a scalable platform for future biomarker discovery efforts. By integrating regularization-based feature selection (e.g., L1-penalized regression) with stability-aware recursive elimination, our approach not only yields concise gene sets with high discriminative power but also enhances interpretability, ensuring that identified markers align with known pathogenic pathways such as oxidative stress, inflammatory signaling, and neurodegeneration. The deployment of multilayer perceptron classifiers further demonstrates how non-linear modeling can capture complex gene–gene interactions and cellular signatures, achieving robust disease-state stratification across diverse retinal cell types. Importantly, machine learning not only validates existing biological knowledge but also uncovers novel cellular behaviors and candidate gene markers such as ectopic gene expression in infiltrating immune cells that warrant targeted experimental validation. Furthermore, the integration of these computational strategies with emerging modalities (e.g., spatial transcriptomics, multi-omics integration, and longitudinal sampling) promises to refine our mechanistic understanding of diabetic retinopathy and other complex diseases, ultimately accelerating the translation of single-cell discoveries into predictive diagnostics and precision therapeutics.

Despite the comprehensive insights gained, this study was subject to several limitations inherent to both single-cell transcriptomic methodologies and machine learning applications. Firstly, while we achieved high classification accuracy, the model’s generalizability to other ethnic populations or clinical subtypes of DR remains to be tested. Secondly, scRNA-seq data are characterized by high sparsity and technical noise, including dropout events where transcripts are undetected despite being present. These artifacts can obscure true biological signals and complicate downstream analyses. Thirdly, although our machine learning models, such as the multilayer perceptron classifier, demonstrated high accuracy in classifying disease states, the “black box” nature of machine learning models poses challenges for interpretability, making it difficult to elucidate the biological underpinnings of the predictive gene features. Lastly, given that the 20 samples analyzed were obtained from the retinas of both eyes of 13 donors, uncertain factors such as interocular differences, disease duration, and inter-individual variability may introduce errors into the analysis. While batch effect correction methods were employed, residual confounding factors related to sample processing and sequencing platforms may persist, potentially influencing the observed gene expression patterns. Future studies integrating multi-omics approaches and larger, more diverse cohorts are warranted to validate and extend these findings.

In conclusion, this study provided a detailed molecular landscape of the Chinese diabetic human retina at single-cell resolution and demonstrated the power of machine learning in identifying reproducible disease markers. By linking specific gene expression changes to retinal cell types and disease states, we offered novel insights into the cellular pathology of DR and lay the groundwork for precision diagnostics and targeted therapeutic interventions. Future directions will include functional validation of candidate genes, integration with spatial and multi-omics data, and deployment in prospective clinical studies to advance personalized management of diabetic retinal diseases.

## Statements and Declarations

### Authors’ Contributions

SL drafted the paper and performed the analysis. LY, YT, QP, TC contributed to data formatting and correction. YY, JL, YZ, YS, QY, ZL, LC, GM, and RR provided comments on the paper. FL, and WM organized the project and provided comments.

## Acknowledgment

This study was mainly funded by the Pioneer and Leading Goose R&D Program of Zhejiang Province 2023 with reference number 2023C04049 and Ningbo International Collaboration Program 2023 with reference number 2023H025. Additionally, this work was supported by the National Natural Science Foundation of China with reference number 82201227 and the Natural Science Foundation of Guangdong Province, China with reference number 2023A1515011225.

## Consent to Publish

All authors have consent for publication.

## Conflict of Interest

The authors have no conflicts of interest to disclose.

## Ethics approval

This study was approved by the Ethics Committee of the University of Nottingham Ningbo China; The Lihuili Hospital affiliated with Ningbo University; The Affiliated Ningbo Eye Hospital of Wenzhou Medical University.

---

Supplementary Table 1: The important gene lists of each human retinal cell type and the consolidated gene list that contains important genes from all retinal cell types after deduplication.

## Notes

### Competing Interest Statement

The authors have declared no competing interest.

## References

1. Leasher J L, Bourne R R A, Flaxman S R, et al. Global estimates on the number of people blind or visually impaired by diabetic retinopathy: a meta-analysis from 1990 to 2010. Diabetes care, 2016, 39(9): 1643–1649.

2. Kang Q, Yang C. Oxidative stress and diabetic retinopathy: Molecular mechanisms, pathogenetic role and therapeutic implications. Redox biology, 2020, 37: 101799.

3. Tang J, Kern T S. Inflammation in diabetic retinopathy. Progress in retinal and eye research, 2011, 30(5): 343–358.

4. Seo H, Park S J, Song M. Diabetic Retinopathy (DR): Mechanisms, Current Therapies, and Emerging Strategies. Cells, 2025, 14(5): 376.

5. Wei L, Sun X, Fan C, et al. The pathophysiological mechanisms underlying diabetic retinopathy. Frontiers in Cell and Developmental Biology, 2022, 10: 963615.

6. Yang T, Zhang N, Yang N. Single-cell sequencing in diabetic retinopathy: progress and prospects. Journal of Translational Medicine, 2025, 23(1): 49.

7. Zou K, Li X, Ren B, et al. Single-cell analysis identifies MKI67+ microglia as drivers of neovascularization in proliferative diabetic retinopathy. Journal of Translational Medicine, 2025, 23(1): 310.

8. Wang J, Sun H, Mou L, et al. Unveiling the molecular complexity of proliferative diabetic retinopathy through scRNA-seq, AlphaFold 2, and machine learning. Frontiers in Endocrinology, 2024, 15: 1382896.

9. McGinnis C S, Murrow L M, Gartner Z J. DoubletFinder: doublet detection in single-cell RNA sequencing data using artificial nearest neighbors. Cell systems, 2019, 8(4): 329–337. e4.

10. Polański K et al. BBKNN: fast batch alignment of single cell transcriptomes. Bioinformatics, 2020, 36(3): 964–965.

11. Wolf F A, Angerer P, Theis F J. SCANPY: large-scale single-cell gene expression data analysis. Genome biology, 2018, 19: 1–5.

12. Traag V A, Waltman L, Van Eck N J. From Louvain to Leiden: guaranteeing well-connected communities. Scientific reports, 2019, 9(1): 1–12.

13. Nandy P, Unger M, Zechner C, et al. Learning diagnostic signatures from microarray data using L1-regularized logistic regression. Systems Biomedicine, 2013, 1(4): 240–246.

14. Misra P, Yadav A S. Improving the classification accuracy using recursive feature elimination with cross-validation. Int. J. Emerg. Technol, 2020, 11(3): 659–665.

15. Yu H, Samuels D C, Zhao Y, et al. Architectures and accuracy of artificial neural network for disease classification from omics data. BMC genomics, 2019, 20: 1–12.

16. Yon Rhee S et al. Use and misuse of the gene ontology annotations. Nature Reviews Genetics, 2008, 9(7): 509–515.

17. Kanehisa M, Goto S. KEGG: kyoto encyclopedia of genes and genomes. Nucleic acids research, 2000, 28(1): 27–30.

18. Yan W et al. Cell atlas of the human fovea and peripheral retina. Scientific reports, 2020, 10(1): 9802.

19. Mwale P F, Hsieh C T, Yen T L, et al. Chitinase-3-like-1: a multifaceted player in neuroinflammation and degenerative pathologies with therapeutic implications. Molecular Neurodegeneration, 2025, 20(1): 7..

20. Kim R Y, Hoffman A S, Itoh N, et al. Astrocyte CCL2 sustains immune cell infiltration in chronic experimental autoimmune encephalomyelitis. Journal of neuroimmunology, 2014, 274(1-2): 53–61.

21. Hölter P, Kunst S, Wolloscheck T, et al. The retinal clock drives the expression of Kcnv2, a channel essential for visual function and cone survival. Investigative ophthalmology & visual science, 2012, 53(11): 6947–6954.

22. Schulkens I A, Castricum K, Weijers E M, et al. Expression, regulation and function of human metallothioneins in endothelial cells. Journal of vascular research, 2014, 51(3): 231–238.

23. Liu X, Secombe J. The histone demethylase KDM5 activates gene expression by recognizing chromatin context through its PHD reader motif. Cell reports, 2015, 13(10): 2219–2231..

24. Sánchez-Ceinos J, Hussain S, Khan A W, et al. Repressive H3K27me3 drives hyperglycemia-induced oxidative and inflammatory transcriptional programs in human endothelium. Cardiovascular Diabetology, 2024, 23(1): 122.

25. J Rengarajan S, Derks J, Bellott D W, et al. Post-transcriptional cross-and auto-regulation buffer expression of the human RNA helicases DDX3X and DDX3Y. Genome Research, 2025, 35(1): 20–30.

26. Pan X, Tan X, McDonald J, et al. Chemokines in diabetic eye disease. Diabetology & Metabolic Syndrome, 2024, 16(1): 115.

27. Dong K, Chen W, Pan X, et al. FCER1G positively relates to macrophage infiltration in clear cell renal cell carcinoma and contributes to unfavorable prognosis by regulating tumor immunity. BMC cancer, 2022, 22(1): 140.

28. Pei X M, Tam B T, Sin T K, et al. S100A8 and S100A9 are associated with doxorubicin-induced cardiotoxicity in the heart of diabetic mice. Frontiers in physiology, 2016, 7: 334.

29. Vieira S M, Monteiro M B, Marques T, et al. Association of genetic variants in the promoter region of genes encoding p22phox (CYBA) and glutamate cysteine ligase catalytic subunit (GCLC) and renal disease in patients with type 1 diabetes mellitus. BMC Medical Genetics, 2011, 12: 1–6.

30. Crouzier L, Diez C, Richard E M, et al. Loss of Pde6a induces rod outer segment shrinkage and visual alterations in pde6aQ70X mutant zebrafish, a relevant model of retinal dystrophy. Frontiers in Cell and Developmental Biology, 2021, 9: 675517.

31. Mosley A L, Ozcan S. Glucose regulates insulin gene transcription by hyperacetylation of histone h4. Journal of Biological Chemistry, 2003, 278(22): 19660–19666.

32. Fan X, Xu M, Chen X, et al. Proteomic profiling and correlations with clinical features reveal biomarkers indicative of diabetic retinopathy with diabetic kidney disease. Frontiers in Endocrinology, 2022, 13: 1001391.

33. Liukkonen M, Heloterä H, Siintamo L, et al. Oxidative Stress and Inflammation-Related mRNAs Are Elevated in Serum of a Finnish Wet AMD Cohort. Investigative Ophthalmology & Visual Science, 2024, 65(13): 30–30.

34. Luo R, Li L, Xiao F, et al. LncRNA FLG-AS1 mitigates diabetic retinopathy by regulating retinal epithelial cell inflammation, oxidative stress, and apoptosis via miR-380-3p/SOCS6 axis. Inflammation, 2022, 45(5): 1936–1949.

35. Roumeliotis A, Roumeliotis S, Tsetsos F, et al. Oxidative stress genes in diabetes mellitus type 2: association with diabetic kidney disease. Oxidative Medicine and Cellular Longevity, 2021, 2021(1): 2531062.

36. Jan Y H, Lai T C, Yang C J, et al. Adenylate kinase 4 modulates oxidative stress and stabilizes HIF-1α to drive lung adenocarcinoma metastasis. Journal of Hematology & Oncology, 2019, 12: 1–14.

37. Ma P, Zhang P, Chen S, et al. Immune cell landscape of patients with diabetic macular edema by single-cell RNA analysis. Frontiers in pharmacology, 2021, 12: 754933.

38. Wang J, Lin J, Schlotterer A, et al. CD74 indicates microglial activation in experimental diabetic retinopathy and exogenous methylglyoxal mimics the response in normoglycemic retina. Acta diabetologica, 2014, 51: 813–821.

39. Wang X, Wang T, Kaneko S, et al. Photoreceptors inhibit pathological retinal angiogenesis through transcriptional regulation of Adam17 via c-Fos. Angiogenesis, 2024, 27(3): 379–395.

40. Li Z, Li Y, Tian Y, et al. Pan-cancer analysis identifies the correlations of Thymosin Beta 10 with predicting prognosis and immunotherapy response. Frontiers in Immunology, 2023, 14: 1170539.

41. Matet A, Jaworski T, Bousquet E, et al. Lipocalin 2 as a potential systemic biomarker for central serous chorioretinopathy. Scientific Reports, 2020, 10(1): 20175.

42. Gnana-Prakasam J P, Martin P M, Mysona B A, et al. Hepcidin expression in mouse retina and its regulation via lipopolysaccharide/Toll-like receptor-4 pathway independent of Hfe. Biochemical Journal, 2008, 411(1): 79–88.

43. Liu J, Li J, Tang Y, et al. Transcriptome analysis combined with Mendelian randomization screening for biomarkers causally associated with diabetic retinopathy. Frontiers in Endocrinology, 2024, 15: 1410066.

44. Deng S, Zhou H, Xiong R, et al. Over-expression of genes and proteins of ubiquitin specific peptidases (USPs) and proteasome subunits (PSs) in breast cancer tissue observed by the methods of RFDD-PCR and proteomics. Breast cancer research and treatment, 2007, 104: 21–30.

45. Sharma S. Interleukin-6 trans-signaling: a pathway with therapeutic potential for diabetic retinopathy. Frontiers in physiology, 2021, 12: 689429..

46. Huang Y, Peng J, Liang Q. Identification of key ferroptosis genes in diabetic retinopathy based on bioinformatics analysis. Plos one, 2023, 18(1): e0280548.

47. Yang L, Li H, Yu T, et al. Polymorphisms in metallothionein-1 and-2 genes associated with the risk of type 2 diabetes mellitus and its complications. American Journal of Physiology-Endocrinology and Metabolism, 2008, 294(5): E987–E992..

48. Resnikoff H A, Miller C G, Schwarzbauer J E. Implications of fibrotic extracellular matrix in diabetic retinopathy. Experimental Biology and Medicine, 2022, 247(13): 1093–1102.

49. Guo X, Yu X, Zhang Y, et al. A novel glycolysis-related signature for predicting the prognosis and immune infiltration of uveal melanoma. Ophthalmic Research, 2023, 66(1): 692–705.

50. Chen Q, Li F, Gao Y, et al. Identification of FGF13 as a potential biomarker and target for diagnosis of impaired glucose tolerance. International Journal of Molecular Sciences, 2023, 24(2): 1807

51. Zhu X, Chen Z, Wang L, et al. Direct conversion of human umbilical cord mesenchymal stem cells into retinal pigment epithelial cells for treatment of retinal degeneration. Cell death & disease, 2022, 13(9): 785.

52. Zhang X, Zhang F, Xu X. Single-cell RNA sequencing in exploring the pathogenesis of diabetic retinopathy. Clinical and Translational Medicine, 2024, 14(7): e1751.

53. Kohno H, Maeda T, Perusek L, et al. CCL3 production by microglial cells modulates disease severity in murine models of retinal degeneration. The Journal of Immunology, 2014, 192(8): 3816–3827.

54. Del Dotto V, Musiani F, Baracca A, et al. Variants in human ATP synthase mitochondrial genes: biochemical dysfunctions, Associated Diseases, and Therapies. International Journal of Molecular Sciences, 2024, 25(4): 2239.

55. Nhieu J, Lin Y L, Wei L N. CRABP1 in non-canonical activities of retinoic acid in health and diseases. Nutrients, 2022, 14(7): 1528.

56. Lahouaoui H, Coutanson C, Cooper H M, et al. Diabetic retinopathy alters light-induced clock gene expression and dopamine levels in the mouse retina. Molecular vision, 2016, 22: 959.

57. Wang Y, Yang C, Gu Q, et al. KLF4 promotes angiogenesis by activating VEGF signaling in human retinal microvascular endothelial cells. PloS one, 2015, 10(6): e0130341.

58. Taranova O V, Magness S T, Fagan B M, et al. SOX2 is a dose-dependent regulator of retinal neural progenitor competence. Genes & development, 2006, 20(9): 1187–1202.

59. Yao Y, Du Jiang P, Chao B N, et al. GIMAP6 regulates autophagy, immune competence, and inflammation in mice and humans. Journal of Experimental Medicine, 2022, 219(6).

60. Gan D M, Zhang P P, Zhang J P, et al. KISS1/KISS1R mediates Sertoli cell apoptosis via the PI3K/AKT signalling pathway in a high-glucose environment. Molecular Medicine Reports, 2021, 23(6): 477.

61. Wang C, Yu Y, Chang Q, et al. Hypermethylation of GSTM5 and its effect on oxidation in myelodysplastic syndrome. iLABMED, 2024, 2(2): 108–124.

62. Yoo H S, Cockrum M A, Napoli J L. Cyp26a1 supports postnatal retinoic acid homeostasis and glucoregulatory control. Journal of Biological Chemistry, 2023, 299(5).

63. Han P, Chang C P. Long non-coding RNA and chromatin remodeling. RNA biology, 2015, 12(10): 1094–1098.

64. Riaz B, Islam S M S, Ryu H M, et al. CD83 regulates the immune responses in inflammatory disorders. International Journal of Molecular Sciences, 2023, 24(3): 2831.

65. Souri Z, Ahmadieh H. Exploring the Connection Between HLA Class I and Class II Genotypes and Diabetic Retinopathy: A Comprehensive Review of Experimental Evidence. Experimental Eye Research, 2024: 110112.

66. Cheng C L, Molday R S. Changes in gene expression associated with retinal degeneration in the rd3 mouse. Molecular vision, 2013, 19: 955.

67. Mears A J, Kondo M, Swain P K, et al. Nrl is required for rod photoreceptor development. Nature genetics, 2001, 29(4): 447–452.

68. You Z P, Zhang Y L, Li B Y, et al. Bioinformatics analysis of weighted genes in diabetic retinopathy. Investigative Ophthalmology & Visual Science, 2018, 59(13): 5558–5563.

69. Roy S, Kern T S, Song B, et al. Mechanistic insights into pathological changes in the diabetic retina: implications for targeting diabetic retinopathy. The American journal of pathology, 2017, 187(1): 9–19.

